# De novo functional discovery of peptide-MHC restricted CARs from recombinase-constructed large-diversity monoclonal T cell libraries

**DOI:** 10.1101/2024.11.27.625413

**Authors:** Zening Wang, Abhijit Sarkar, Xin Ge

## Abstract

Chimeric antigen receptors (CAR) that mimic T cell receptors (TCR) on eliciting peptide-major histocompatibility complex (pMHC) specific T cell responses hold great promise in the development of immunotherapies against solid tumors, infections, and autoimmune diseases. However, broad applications of TCR-mimic (TCRm) CARs are hindered to date largely due to lack of a facile approach for the effective isolation of TCRm CARs. Here, we establish a highly efficient process for *de novo* discovery of TCRm CARs from human naïve antibody repertories by combining recombinase-mediated large-diversity monoclonal library construction with T cell activation-based positive and negative screenings. Panels of highly functional TCRm CARs with peptide-specific recognition, minimal cross-reactivity, and low tonic signaling were rapidly identified towards MHC-restricted intracellular tumor-associated antigens MAGE-A3, NY-ESO-1, and MART-1. Transduced TCRm CAR-T cells exhibited pMHC-specific functional avidity, potent cytokine release, and efficacious and persistent cytotoxicity. The developed approach could be used to generate safe and potent immunotherapies targeting MHC-restricted antigens.

## INTRODUCTION

The recognition of peptide-major histocompatibility complex (pMHC) molecules by T cells is a key feature of adaptive immunity and provides the basis for the development of immunotherapies against malignancies, infections, and autoimmune diseases. As natural ligands of pMHC, T cell receptors (TCR) have been exploited to develop adoptive cell therapies, in which the promise of TCR gene transfer has been demonstrated especially in the field of cancer immunotherapy [Klebanoff *et al*., 2023; Baulu *et al*., 2023]. However, one major challenge for TCR-engineered T (TCR-T) cell therapy is TCR mispairing and the associated neoactivities [Bendle *et al*., 2010; van Loenen *et al*., 2010]. Although considerable efforts have been made to address this issue [Govers *et al*., 2010; Bunse *et al*., 2014; Bethune *et al*., 2016], achieving complete circumvention of TCR mispairing remains difficult. In this context, an appealing alternative is to develop chimeric antigen receptor (CAR)-T cells which utilizes an antibody fragment (*e.g.*, scFv) as the pMHC-recognizing module and possesses its own T cell activation signaling orthogonal to that of the endogenous TCR [Akatsuka, 2020; Poorebrahim *et al*., 2021; Li *et al*., 2022]. Avoiding endogenous TCR knockout, the manufacturing process of CAR-T cells is facile for standardization [Ceja *et al*., 2024]. Additionally, uses of pMHC targeting TCR mimic (TCRm) antibodies are more versatile, such as the development of bispecific T cell engagers (BiTE) [Fenis *et al*., 2024]. While lacking the assistance given by CD8 coreceptor and the FG loop of TCRβ constant domain, both enhancing antigen sensitivity of TCRs [Des *et al*., 2015; Clement *et al*., 2021], it has been proved that the inherent high affinities and slow dissociation rates of antibodies can render a prolonged pMHC recognition needed for effective T cell stimulation [Banik *et al*., 2021]. Indeed, TCRm antibodies that replicate the canonical footprint on pMHC and the natural binding orientation of TCRs have been identified [Holland *et al*., 2020], and functional similarities between TCRs and TCRm CARs on eliciting pMHC-restricted T cell activation have been demonstrated [Wang *et al*., 2021]. To date, TCRm CARs and T cell engagers are emerging as attractive therapeutics towards neoantigens [Hsiue *et al*., 2021; Douglass *et al*., 2021; Dao *et al*., 2022; Yarmarkovich *et al*., 2023], tumor-associated antigens (TAAs) [Raskin *et al*., 2021; Klatt *et al*., 2022; Liu *et al*., 2022; Wang L *et al*., 2024], HPV oncogenes [Duan *et al*., 2024], HIV peptides [Sengupta *et al*., 2022], and T1D autoantigens [Zhang *et al*., 2019].

Despite the importance of TCRm CARs, their broad applications have been largely hindered because current monoclonal antibody (mAb) discovery approaches are incompetent for functional CAR isolation. Unlike mAbs as soluble therapies, CARs act on T cell surface, and their functions are carried out by the cells displaying them. Upon pMHC ligation and the formation of immunological synapse, T cell activation is initiated through mechanotransduction (cytoskeletal rearrangement), which triggers a cascade of signaling events including ITAM phosphorylation, transcriptional regulation, and T cell proliferation [Salter *et al*., 2021]. These complex cellular responses extend far beyond simple antigen-antibody interactions that underpin most mAb discovery strategies to date. The miscorrelation between pMHC binding and T cell activation is evidenced by the observation that although pMHC specific binders can be readily isolated through methods like phage panning, once converted to CARs, they often fail to effectively stimulate pMHC-specific T cell effector function. To mitigate this challenge, elegant library designs focusing on peptide-contacting residues have been employed to engineer TCRm Abs of canonical docking topologies for improved affinity [Stewart-Jones *et al*., 2009] or altered specificity toward new MHC-restricted peptide antigens [Yang *et al*., 2023]. However, efficient *de novo* discovery of functional TCRm CARs has not been fulfilled. Series of mechanistic studies have revealed three desired biophysical properties essential for effective and specific T cell activation – a moderate affinity [Maus *et al*., 2016], a broad and energy balanced epitope [Holland *et al*., 2020], and a prolonged lifetime of the interaction via slow dissociation off-rates [Hebeisen *et al*., 2015] or the formation of catch bonds [Sibener *et al*., 2018; Zhu *et al*., 2019; Wang 2020; Choi *et al*., 2023]. These insights suggest that for the efficient isolation of TCRm CARs, it is necessary to utilize T cells and reconstitute their activation process within, *i.e.*, construction of large-diversity CAR libraries, coculture with antigen presentation cells (APCs), and high-throughput screening for T cell activation events. Furthermore, given the adverse effects associated with peptide independent cytotoxicity [Linette *et al*., 2013; Morgan *et al*., 2013] and the diminished antitumor efficacy caused by early T cell exhaustion [Long *et al*., 2015], strategies to eliminate clones of cross-reactivity and spontaneous activation are also critically needed.

Carrying the signaling machinery and transcriptional network of primary T cells, immortalized human T lymphocytes including Jurkat or SKW-3 have been engineered for functional CAR or TCR screening, *e.g.*, by using NFAT-EGFP as a reporter or monitoring activation markers such as CD69 [Rydzek *et al*., 2019; Di Roberto *et al*., 2020; Spindler *et al*., 2020; Fahad *et al*., 2022; Vazquez-Lombardi *et al*., 2022; Zhao *et al*., 2022; Moravec *et al*., 2024]. However, current methods of combinatorial library construction in immortalized T cells are suboptimal, representing a significant bottleneck in the rapid discovery of effective TCRs or TCRm CARs. Viral vectors are commonly used for gene delivery to T cells, but their characteristic of random genomic integration leads to polyclonality (multiple clones per cells) and transgene transcription variation, both of which cause false events and complicate the identification of candidate clones. To achieve single clone per cell, low multiplicity of infection (MOI) is often applied, which nonetheless compromises transduction efficiency while monoclonality is still not guaranteed. In addition, the labor-intensive viral packaging process plausibly introduces biases among library members, potentially causing loss of rare yet valuable clones. Recent advances in CRISPR/Cas9 genome editing have enabled targeted transgene integration in Jurkat cells such as at TRAC and TCRβ loci [Di Roberto *et al*., 2020; Vazquez-Lombardi *et al*., 2022]. Using this approach, site-directed mutagenesis libraries focusing on CDR-H3 and CDR3β regions have been constructed, leading to the isolation of CAR and TCR variants with enhanced potency and/or specificity. Though suitable for CDR fine-tuning, the achieved diversities of ∼10^5^ variants so far are orders of magnitude smaller than the desired library size, *i.e.*, 10^7^ or more, needed for full-length TCR and CAR discovery and engineering. In addition, the erroneous nature of nuclease-initiated DNA repair process often results in indels and unintended mutants [Allen *et al*., 2018; Boel *et al*., 2018], and transgene biallelicity has been observed as well [Parthiban *et al*., 2019].

Catalyzing genomic integration of large transgenes with strict accuracy, site-specific recombinases overcome the critical limitations of earlier methods and have been implemented for multiplex genetic assays and directed evolution in mammalian cells [Segaliny *et al*., 2023; Maricque *et al*., 2019; Cao *et al*., 2021; Chen C *et al*., 2023; Wang Z *et al*., 2024a; Wang Z *et al*., 2024b]. Particularly, advancement by Durrant *et al*. has identified large serine recombinases (LSR) exhibiting remarkable efficiency with minimal off-target integration in human cells [Durrant *et al*., 2023]. In this study, we apply LSR Pa01 in Jurkat cells to achieve ∼40% genomic integration efficiency of multi-kilobase DNA payloads, including TCR and CAR transgenes, in a monoclonal, site-specific, and cell viability-friendly manner. This breakthrough enables facile construction of large-diversity human naïve antibody repertoires derived CAR libraries of 2.0×10^7^ clones and efficient screening for functional TCRm CARs. We present data on the rapid identification of panels of CAR clones that specifically and effectively elicit T cell responses to HLA-A*02:01 restricted peptides of TAAs including MAGE-A3, NY-ESO-1, and MART-1. We also demonstrate the minimal cross-reactivity and low tonic signaling of isolated CAR clones and their specific and potent T cell cytotoxicity against the respective tumor cells. We believe the technology described here is widely applicable for targeting various pMHC molecules of biomedical importance and can substantially advance the development of diverse immunotherapy modalities including CAR-T, TCR-T, and T cell engagers.

## RESULTS

### Recombinase-mediated high efficiency transgene integration in Jurkat cells

Our initial tests of LSRs in polyclonal Jurkat cells indicated that recombinase Pa01 rendered 27±1.8% transgene integration efficiency (17-fold higher than Bxb1) without impacting cell viability (>95%) (**Fig. S1**). To further evaluate Pa01, we established monoclonal Jurkat cell lines harboring single genomic landing pads encoding an attB-Pa01-2A-EGFP cassette downstream of the pEF1a promoter (**Fig. 1a**). Jurkat-Pa01 clones E11 and B4 exhibiting stable and high EGFP expression were isolated (**Fig. S2**), and their single landing pads in genomes were confirmed by genotyping (**Fig. S3**). On the donor plasmids, a promoterless GOI is sandwiched between a Pa01 recognition site and a transcription terminator, and thus a successful genomic integration would lead to GOI expression and concomitant loss of Pa01 and EGFP production (**Fig. 1a**). When mCherry was tested as the GOI, on day 5 post electroporation, 33±3% of E11 cells and 31±6% of B4 cells were mCherry^+^. The transgene integration efficiencies increased to 39±3% and 37±6% on day 12 and plateaued at 42±4% and 38±6% on day 19, respectively (**Fig. 1b**). Notably, both the initial background expression of mCherry and residual EGFP signals in the transgene integrated cells diminished over the time. Furthermore, mCherry^+^/EGFP^-^ cell populations clustered on flow cytometry plots, suggesting universal transcriptional levels likely due to the monoclonality and the same genomic context governed by site-specific recombination. Besides stable integration and sustained transgene expression, both engineered Jurkat cell lines maintained high cell yields and viabilities (*e.g.*, > 86% on day 1 with full recoveries on day 2 at ∼94%), exhibiting minimal toxicity from donor DNA electroporation. Similar high integration efficiencies (up to 46% in E11 and 37% in B4) and >95% cell viabilities were observed when using iRFP as the GOI (**Fig. S4**). To further validate monoclonality of Pa01-mediated transgene integration, a mixture of equal molar of mCherry and iRFP donor plasmids was applied for electroporation, and only single positive cell populations were obtained, confirming that Jurkat-Pa01 lines E11 and B4 indeed performed single integration events (**Fig. 1c**).

**Figure 1.**
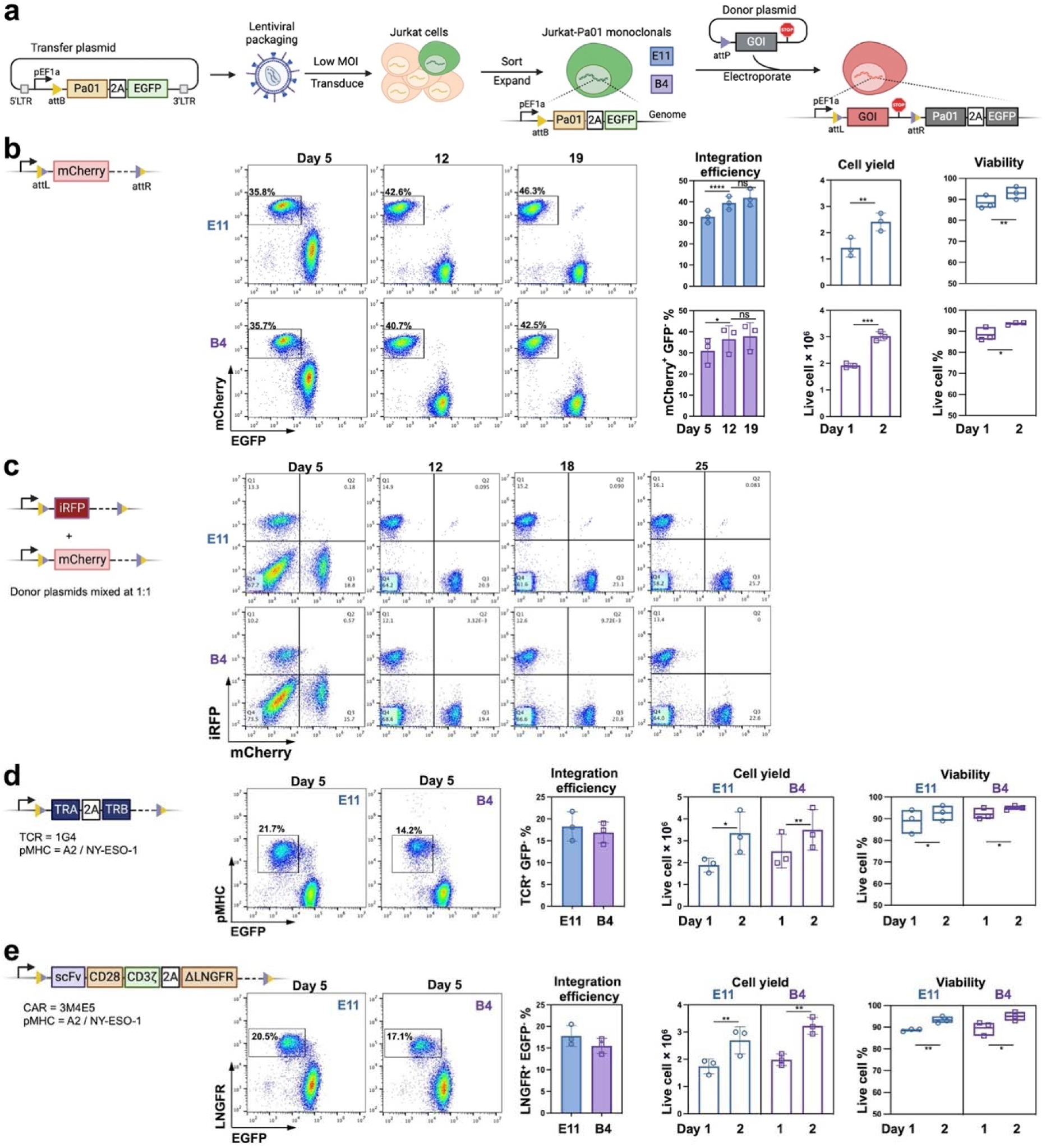
Recombinase-mediated high-efficiency monoclonal transgene integration to Jurkat cell lines. **a,** Schematic. Generation of monoclonal cell lines carrying the attB-Pa01-2A-EGFP landing pad for transgene genomic integration. Donor plasmid encodes a promoter-less attP-GOI cassette followed by a transcription terminator. Upon Pa01-catalyzded recombination, GOI is produced while Pa01-2A-EGFP expression is silenced. **b,** mCherry was tested as the GOI for integration in monoclonal Jurkat-Pa01 cell lines E11 and B4. Two million cells were electroporated with 4 µg of donor plasmid. The integration efficiency was analyzed by flow cytometry on day 5, 12, and 19 post electroporation, and the cell yields and viability were monitored on day 1 and 2 post electroporation. **c,** Validation of monoclonality of transgene integration. Jurkat-Pa01 E11 and B4 cells were electroporated with a mixture of equal molar of mCherry and iRFP donor plasmids, and cells were monitored by flow cytometry till day 25. **d**, TCR, or **e**, CAR was tested as the GOI. Means and s.d. are plotted from three independent experiments. A one-way ANOVA with Turkey’s test statistic was conducted for integration efficiency; a one-tailed t test was conducted for cell yield and viability. ns, not significant; *, p≤ 0.0332; **, p≤ 0.0021; **** p≤ 0.0001.

To evaluate the integration efficiencies of TCR and CAR transgenes, E11 and B4 cells were electroporated with promoterless donor plasmids encoding 1G4 TCR (**Fig. 1d**) or 3M4E5 CAR (**Fig. 1e**), both targeting HLA-A*02:01 restricted peptide NY-ESO-1_157-164_ (SLLMWITQC). Results indicated that 18±3% and 17±2% of 1G4 TCR donor DNA electroporated E11 and B4 cells were NY-ESO-1_157-165_/A2 tetramer positive on day 5 post electroporation (**Fig. 1d**). For 3M4E5 CAR, its transgene integration efficiencies, determined by the LNGFR^+^/EGFP^-^ populations as its construct was fused with a truncated LNGFR via a 2A self-cleaving peptide linker, achieved 18±2% and 15±2% in E11 and B4 cells, respectively (**Fig. 1e**). We also tested the HLA-A*02:01 restricted MART-1_26-35(27L)_ (ELAGIGILTV) specific MA2 CAR [Yang *et al*., 2023], and a high transgene integration efficiency at 17% was similarly obtained in B4 cells (**Fig. S5**). Consistent with mCherry and iRFP donors, high cell viability (*e.g.*, >93% on day 2) and normal cell growth were observed for electroporation with TCR and CAR donor plasmids (**Fig. 1de**). Overall, these results indicated high integration efficiencies of multi-kilobase DNA payloads including TCR and CAR transgenes into the genome of Jurkat cells in a monoclonal, site-specific, and cell viability-friendly manner.

### Validation of high-throughput functional screening for TCRm CAR

The functionalities of TCRs and CARs displayed on Jurkat-Pa01 B4 cells were validated on day 12 post electroporation by both binding assays with pMHC tetramers and activation assays via monitoring the early activation marker CD69 following co-cultures with cells presenting the cognate or control peptide (**Fig. 2a**). Results indicated that 96% of 3M4E5 CAR transgene integrated B4 cells (EGFP^-^) bound to its cognate NY-ESO-1_157-165_/A2 tetramer rather than the irrelevant MART-1_26-35(27L)_/A2 tetramer (**Fig. 2b**). Activation assays further demonstrated that upon stimulation with NY-ESO-1_157-164_ pulsed APCs, CD69 was upregulated among 87% of the 3M4E5 CAR expressing (NY-ESO-1/A2^+^) cells, while this number dropped to 5.7% when MART-1_26-35(27L)_ pulsed APCs were applied (**Fig. 2c**). Such pMHC specific recognition and activation were observed in 1G4 TCR and MA2 CAR transgene integrated B4 cells as well (**Fig. 2bc**, **Fig. S5**), collectively confirming the biological activities of displayed TCR and CAR transgenes in Jurkat-Pa01 cells. Notably, the activated B4 cells formed populations distinct from these of inactivated cells on flow cytometry plots, providing foundation for the development of a high-throughput screening (HTS) approach for the isolation of pMHC specific and functional TCR/CAR clones.

**Figure 2.**
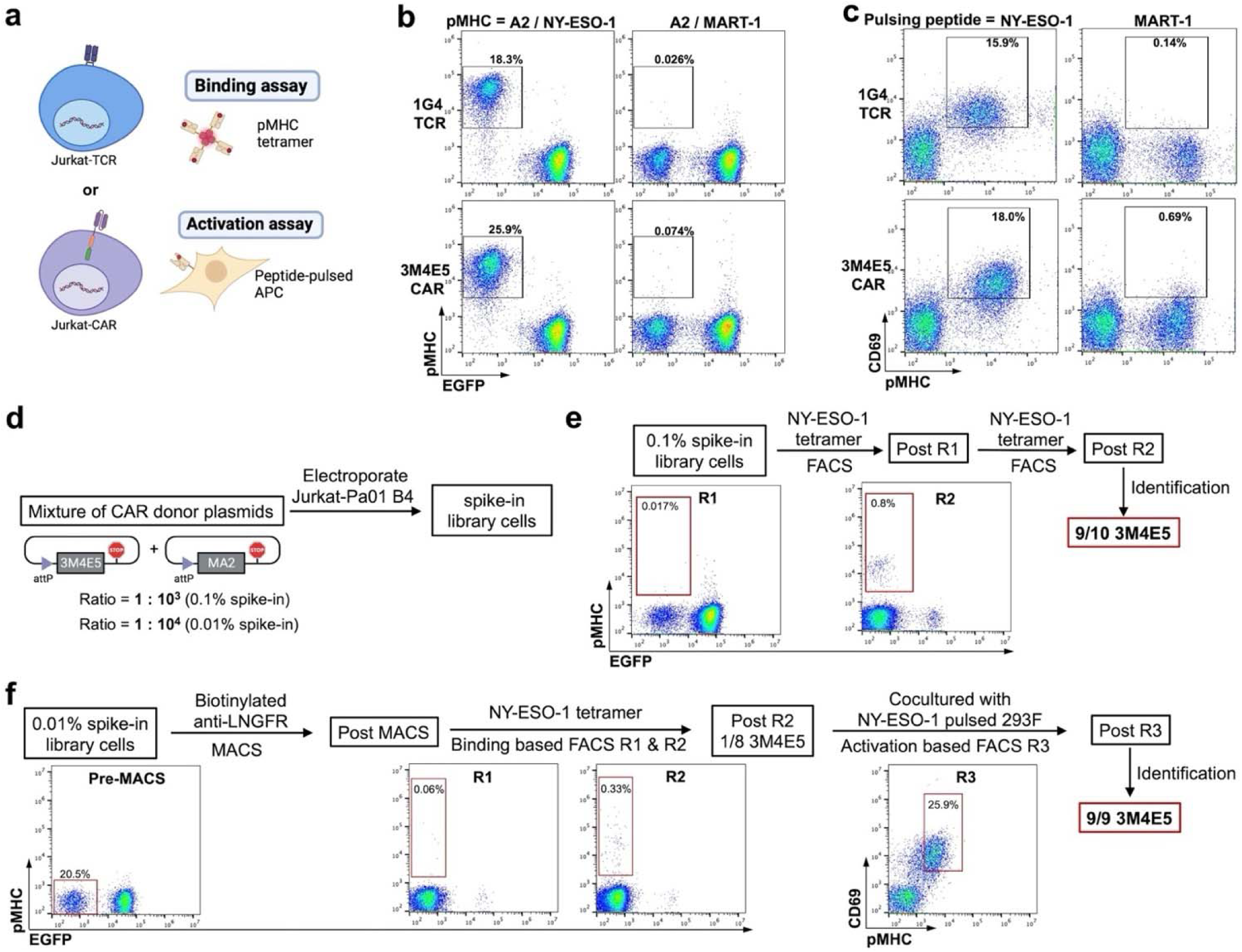
Validation of high-throughput screening of TCRm CAR libraries. **a,** Functional display of TCR and TCR-like CAR on Jurkat-Pa01. B4 cells displaying TCR or CAR were subjected to binding assays with pMHC tetramer and activation assays by co-culture with peptide-pulsed APCs. **b,** Binding assay. Jurkat-Pa01 B4 cells were electroporated with donor plasmids attP-1G4 or attP-3M4E5, stained with cognate A2/NY-ESO-1_157-165_ or irrelevant A2/MART-1_26-35(27L)_, and analyzed by flow cytometry on day 5. **c,** Activation assay. Jurkat cells displaying 1G4 TCR or 3M4E5 CAR were overnight co-cultured with 293F cells pulsed with 10 µM NY-ESO-1 or MART-1 peptide. In the next day, cells were stained with A2/NY-ESO-1 tetramer and upregulation of CD69 was monitored. **d,** Construction of 0.1% and 0.01% spike-in libraries by electroporating 2 and 10 million of Jurkat-Pa01 B4 cells with 4 and 20 µg of a mixture of 3M4E5 and MA2 CAR donor plasmids at a ratio of 1:10^3^ and 1:10^4^, respectively. **e,** Screening 0.1% spike-in library. The library cells were subjected to two rounds of binding based FACS to isolate A2 / NY-ESO-1 pMHC tetramer bound cells. The sorted cells from the second round were directly used for CAR identification including genomic DNA extraction, CAR gene amplification and subcloning, *E. coli* transformation, single colony picking, minipreps, and sanger sequencing. The result showed 9 out of 10 colonies were 3M4E5 CAR. **f,** Screening 0.01% spike-in library. The library cells were first subjected to one round of MACS using biotinylated anti-LNGFR to enrich CAR displayed cells. The enriched cells were expanded and subjected to 2 rounds of binding based FACS to isolate A2 / NY-ESO-1 pMHC tetramer bound cells. After expansion, post R2 cells were used for CAR identification or subjected to one round of activation based FACS after overnight co-culture with NY-ESO-1 peptide pulsed 293F cells. The A2 / NY-ESO-1 pMHC tetramer bound cells with upregulated CD69 were isolated and expanded followed by CAR identification. The results showed 1 out of 8 colonies was 3M4E5 CAR after R2 whereas 9 out 9 were after R3.

To evaluate the suitability of recombinase-constructed libraries for HTS, we conducted control experiments by spiking donor plasmid of 3M4E5 CAR (as the positive clone) into that of MA2 CAR (as the background clone) at a sensitivity frequency of 10^-3^ (0.1%) or 10^-4^ (0.01%) (**Fig. 2d**). Electroporation of 10 million Jurkat-Pa01 B4 cells with the 0.01% spiked donor plasmid mixture resulted in 20.5% EGFP^-^ cell populations, indicating the feasibility to construct a CAR library of >2×10^6^ clones in Jurkat cells. With the 0.1% spike-in library, two rounds of fluorescence-activated cell sorting (FACS) were performed based on binding to NY-ESO-1/A2 pMHC tetramer, and 0.2-0.8% of cells with the highest signals were isolated. Results indicated that pMHC specific cell population enriched from 0.017% in the spike-in library to 0.8% after R1, and among the post-R2 population, Sanger sequencing identified 3M4E5 clone at an abundancy of 9/10 (**Fig. 2e**). For the 0.01% spike-in library, cells were labeled with biotinylated anti-LNGFR and captured by streptavidin beads in magnetic-activated cell sorting (MACS). Results showed that 95% of post-MACS cells were EGFP^-^, indicating successful enrichment of CAR transgene integrated cells (**Fig. 2f**). These post-MACS cells were then subjected to two rounds of binding based FACS, and pMHC^+^ populations increased to 0.33% after R1 and 17% after R2, consistent with the sequencing results on the post-R2 samples (1/8 was 3M4E5). We conducted activation based FACS in R3, via isolation of CD69^+^/pMHC^+^ Jurkat cells following overnight coculture with NY-ESO-1_157-165_ pulsed HLA-A*02:01 positive 293F cells. Flow cytometry analysis suggested that the entire post-R3 population was NY-ESO-1_157-165_/A2 positive, and sequencing results confirmed that 9/9 clones were 3M4E5 CAR (**Fig. 2f**). Overall, spike-in experiments proved the feasibility of HTS for specific and functional TCRm CAR, allowing us to perform *de novo* discovery next.

### Discovery of functional TCRm CARs targeting TAAs from human naïve antibody repertoires

Having demonstrated the effectiveness and sensitivity of our platform, we sought to construct large libraries of human naïve antibodies derived CARs in Jurkat cells and screen for TCRm CAR clones targeting HLA-A*02:01 restricted peptides of three public TAAs: MAGE-A3_112-120_ (KVAELVHFL), NY-ESO-1_157-164_, and MART-1_26-35(27L)_. As well-known cancer-testis antigens (CTAs), MAGE-A3 and NY-ESO-1 are tumor-specific, immunogenic, and uniquely expressed in a wide range of cancer types [Baulu *et al*., 2023; Klebanoff *et al*., 2023; Peri *et al*., 2023]. Despite their promising potentials in cancer immunotherapy, a safe and effective TCR-or CAR-T cell therapy remains to be improved. To discover highly specific and efficacious TCRm CARs, we developed a workflow combining pMHC binding and APC-mediated T cell activation (**Fig. 3a**). Particularly, APCs were pulsed with the cognate and control peptides for positive and negative screenings, respectively, to isolate functional TCRm CARs and remove cross-reactive or self-activated clones. We monitored the upregulation of CD69, a well-defined early marker of T cell activation, as a dual-purpose readout for both CAR-pMHC recognition induced activation and antigen-independent tonic signaling [Chen J *et al*., 2023].

**Figure 3.**
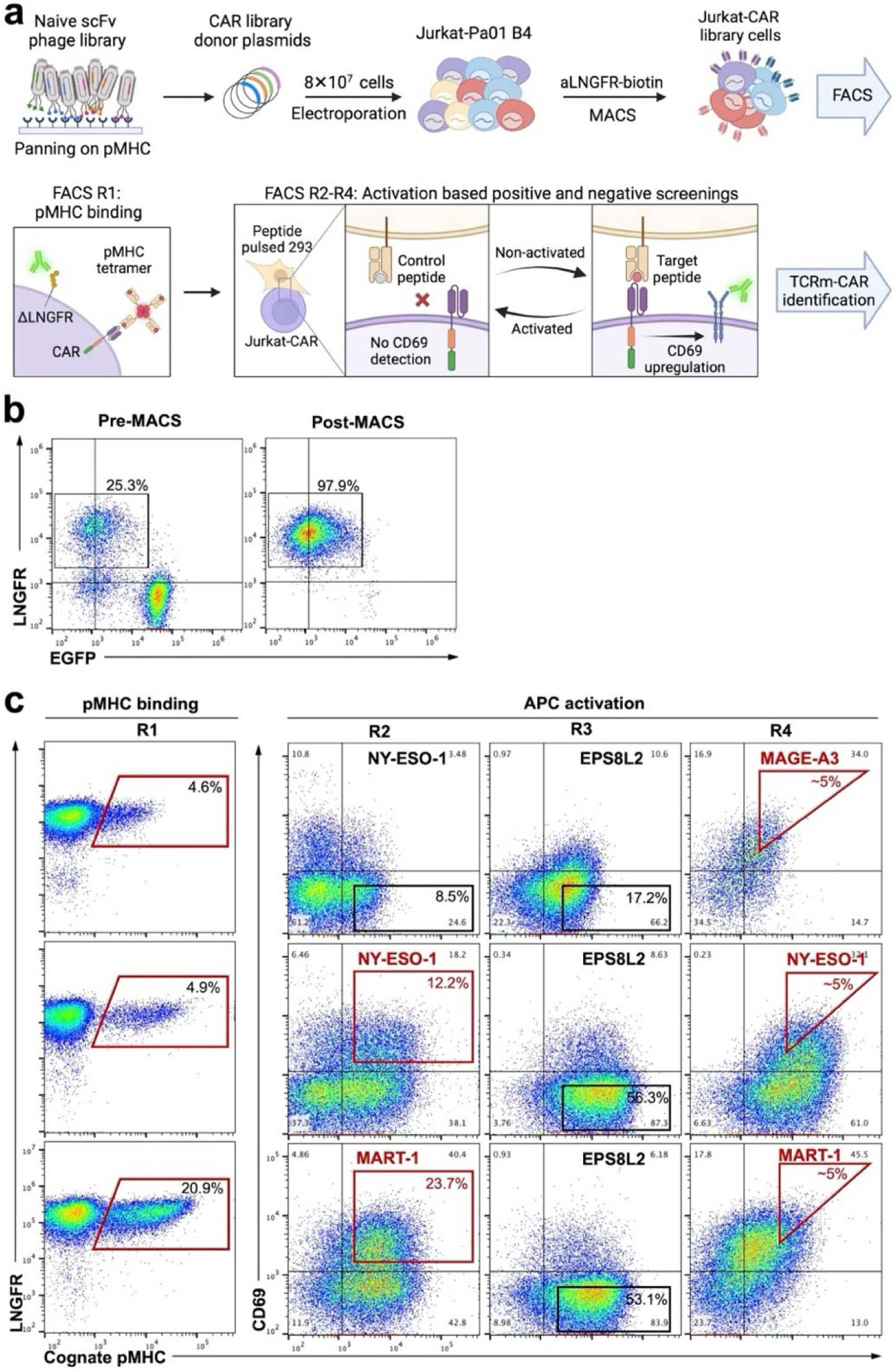
High-throughput screening for specific, functional, and not self-activated TCRm CAR clones. HLA-A*02:01 restricted peptides MAGE-A3_112-120_, NY-ESO-1_157-164_, and MART-1_26-35(27L)_ were used as the targets. **a,** Workflow. Following two rounds of phage panning on each pMHC, down-sized scFv libraries were cloned into CAR donor plasmids, and electroporated to 8×10^7^ Jurkat-Pa01 B4 cells to construct Jurkat-CAR libraries. CAR displayed cells were enriched by MACS and then subjected to 4 rounds of FACS – R1 for pMHC binders, and activation-based screening or counter-screening in R2-R4 by co-culturing with respective APCs pulsed with either the target or irrelevant peptides. **b,** Representative flow cytometry plots showing the HLA-A2/NY-ESO-1_157-164_ targeting Jurkat-CAR library cells before and after MACS. **c,** FACS results. In R2-R4 of activation screening, APC were pulsed with the indicated peptides. After coculture, CAR-T cells were stained by the cognate pMHC and anti-CD69. Target peptides and positive gates are in red; irrelevant peptides and negative gates are in black.

A human naïve scFv library of >10^9^ clones [Ku *et al*., 2021] was firstly downsized by two rounds of phage panning towards each pMHC target. To enrich specific binders, depletion steps using empty MHC and irrelevant pMHC monomers were applied, and 2.0-2.7×10^6^ phages were isolated for each pMHC target. The isolated CAR genes were then subcloned to construct Pa01 attP-CAR library donor plasmids, and 1.2-2.2×10^8^ *E. coli* transformants were obtained per library. Following electroporation of 8.0×10^7^ Jurkat-Pa01 B4 cells and expansion, on day 7, flow cytometry analysis revealed that ∼25% of the transfected CAR library cells were LNGFR^+^ (**Fig. 3b**, pre-MACS), indicating 7.4-and 10-fold diversity coverages over the panned libraries. To remove cells lacking transgene integration or expression, 2.0×10^8^ transfected B4 cells of each library were subjected to MACS with anti-LNGFR coated magnetic beads. 4.0-5.0×10^7^ cells were isolated from each library, and the flow cytometry analysis confirmed that 98% post-MACS cells were LNGFR^+^ (**Fig. 3b**, post-MACS).

To isolate pMHC-specific and APC responsive TCRm CAR clones from these three post-MACS pools, we conducted four rounds of FACS – binding based screening in R1, and activation-based screening / counter-screening in R2-R4 (**Fig. 3a**; gates shown in **Fig. 3c**). In R1, the 4.6-21% of post-MACS cell populations positive to their cognate pMHC tetramers were isolated (**Fig. 3c**, R1; also indicating the successful enrichment of pMHC binders via phage panning). In R2-R4, library cells were co-cultured with either the target or control peptide pulsed 293F cells and assessed by both cognate pMHC binding and CD69 activation marker. Taking NY-ESO-1 library cells in R2 as an example, 56% of the cells were NY-ESO-1_157-165_/A2 positive (**Fig. 3c**, R2), suggesting further enrichment of the binders in R1. However, only one-third of pMHC^+^ cells exhibited CD69 upregulation when exposed to cognate peptide pulsed APCs (functional clones), while 38% of the library cells were pMHC positive but CD69 negative (suggesting affinity but nonfunctional clones). Notably, 6.5% of the cells were negative on binding to the cognate pMHC yet activated in an antigen-independent manner (self-activated clones). All these observations highlighted the critical role of activation-based screening to identify TCRm CARs that not only specifically bind but also functionally respond to the cognate pMHC displayed by APCs. In R2, the top pMHC^+^ and CD69^+^ double positive cells (∼12%) were isolated from NY-ESO-1 library cells, and similar positive screening was applied to MART-1 library cells (**Fig. 3c**, R2). In R3, peptide EPS8L2_339-347_ (SAAELVHFL), to which anti-MAGE-A3 TCRs are cross-reactive [Martin *et al*., 2021], was applied for counter-screenings. Results showed that 7-11% of the cells were CD69^+^ (**Fig. 3c**, R3), indicating either spontaneous or non-specific activation in response to APCs carrying this control/irrelevant peptide (cross-reactive clones). These results underscored the necessity of counter-screenings to eliminate cross-reactive and self-activated CAR clones. Accordingly, pMHC^+^/CD69^-^ cell populations were isolated in R3. And in R4, library cells were co-cultured with APCs carrying the cognate pMHC, and flow cytometry results clearly showed the enrichments of populations that were both bound to the cognate pMHC and stimulated by the cognate APCs (**Fig. 3c**, R4). The top double positive cells (∼5%) were isolated in R4, representing the most promising pMHC-specific and functional TCRm-CAR candidates for further screening and characterizations.

### Identification of functional TCRm CAR clones

To identify isolated TCRm-CAR clones, genomic DNA was extracted from the post-R4 cells and the amplified scFv genes were subcloned *en masse* into the lentiviral plasmid designed for CAR display and mCherry co-expression (**Fig. 4a**). Out of 60-100 randomly picked colonies, Sanger sequencing identified 9, 34, and 42 CDR unique scFv clones for HLA-A*02:01 restricted MAGE-A3_112-120_, NY-ESO-1_157-164_, and MART-1_26-35(27L)_ peptides, respectively (denoted as A3, NY, and MR TCRm CAR clones). Antibody germline analysis revealed diverse IGHV and IGLV/IGKV gene usage among the isolated TCRm CARs (**Fig. S6**). While distinct preferences were observed, interestingly IGHV1-69 and IGKV1-39 were dominant across targets. To validate specificity and functionality of identified TCRm CARs, we transduced the packaged lentivirus into a Jurkat reporter cell line carrying an NFAT-EGFP cassette and co-cultured them with APCs presenting the cognate or control peptides (**Fig. 4a**). The levels of T cell activation were determined as the relative percentages of EGFP^+^ cells normalized to CAR displaying mCherry^+^ cells (**Fig. S7**). As results summarized in **Fig. 4b-d**, astonishingly 7 out of 9 A3, 31 out of 34 NY, and 41 out of 42 MR TCRm CAR clones exhibited significant T cell activation. This high success rate indicates the effectiveness of our naïve TCRm CAR discovery approach, presumably due to the combination of large-diversity monoclonal CAR library construction and activation-based screenings. Specificity testing showed that the majority of isolated NY and MR TCRm CAR clones exhibited negligible T cell activation when exposed to APCs pulsed with the irrelevant peptide EPS8L2_339-347_, with only a few exceptions (NY-38, MR-32, and MR-74), indicating efficient counter-screenings (**Fig. 4c-d**). Similarly, all seven identified functional A3 TCRm CAR clones demonstrated 4.9-to 11-fold differences in T cell activation when co-cultured with cognate peptide versus negative peptide pulsed 293F cells (**Fig. 4d**). Overall, panels of specific and functional TCRm CAR clones targeting three TAAs were successfully discovered.

**Figure 4.**
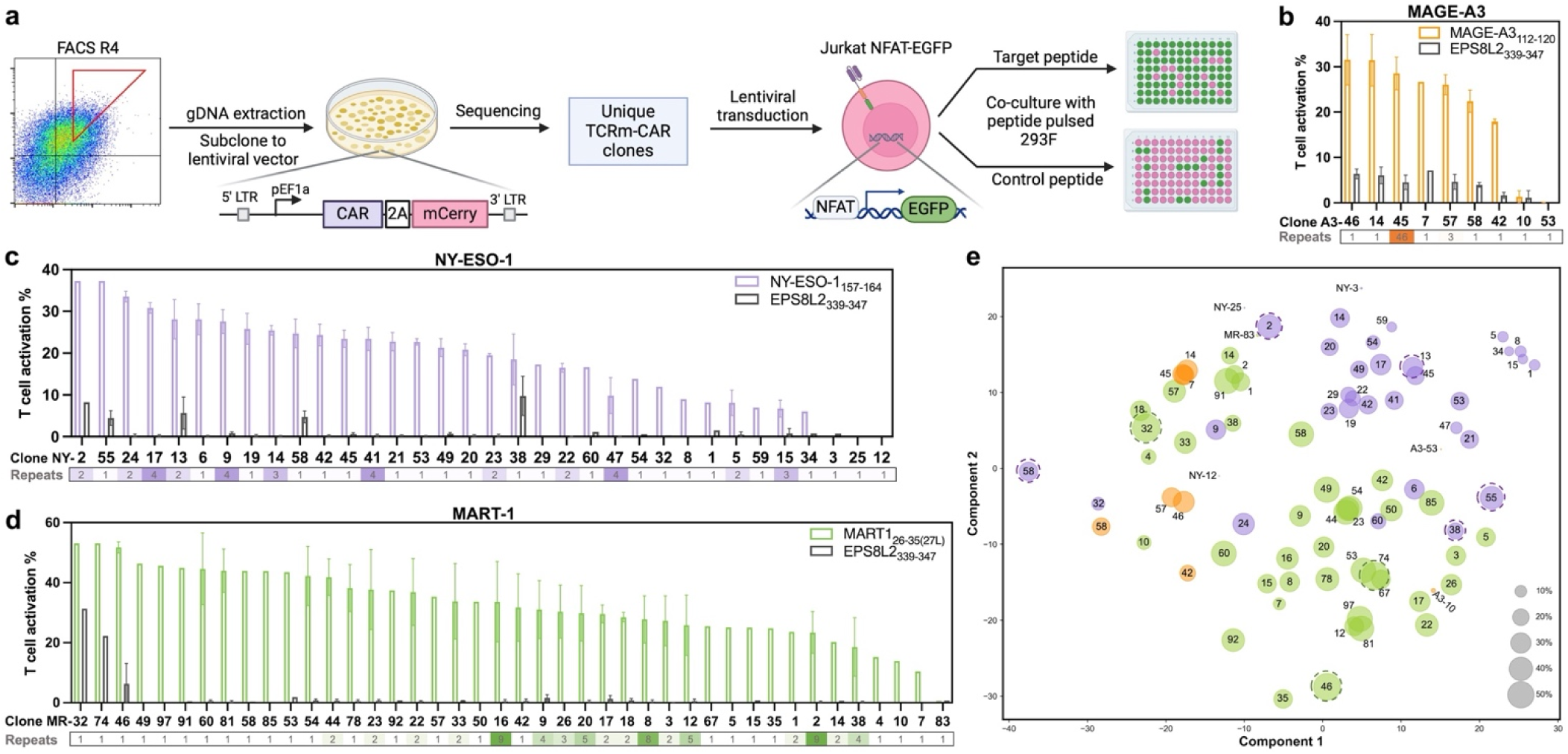
TCRm-CAR identification and specificity screening. **a**, Schematic. Genomic DNA was extracted from post-R4 cells, and the selected TCRm-CAR genes were subcloned in frame with 2A-mCherry to lentiviral vectors for unique clone identification and transduction of Jurkat NFAT-EGFP cells. TCRm-CAR transduced Jurkat NFAT-EGFP cells were cocultured with either target or control peptide pulsed 293F cells, and mCherry and EGFP signals were monitored as indications of CAR display and T cell activation, respectively. **b**, MAGE-A3, **c**, NY-ESO-1, and **d**, MART-1 targeting TCRm-CAR clones. Clones were ranked by T cell activation, which was calculated as the percentage of EGFP positive cells normalized to CAR positive cells. Abundancies of each identified clones are shown at bottom. EPS8L2 peptide pulsed 293F cells were used to test the cross-reactivity. **e,** CDR sequence distance analysis. Distance measurements between any two CAR clones were conducted by calculation of weighted mismatch distance between the amino acid sequences of CDRs using the BLOSUM62 substitution matrix. The results of clone clustering were visualized by multidimensional scaling (MDS) – the more similar the CDRs of the clones, the closer on the MDS plot. Each disk represents a unique clone (A3, orange; NY, purple; MR, green), and its size indicates the strength of the CAR clone in T cell activation assays. Clones having significant cross reactivities are dash circled.

It is remarkable that within human naïve antibody repertoire, we found a vast array of clones capable of targeting MHC-presented intracellular TAAs – a function typically associated with TCRs. Inspired by the observation that TCRs recognizing the same pMHC often share conserved sequence features [Dash *et al*., 2017], we wonder whether our identified TCRm CAR clones exhibit a similar characteristic. A distance matrix between any two CAR clones was computed based on similarity-weighted mismatch of their CDR sequences. Visualization of the distance analysis through multidimensional scaling (MDS) revealed that TCRm CAR clones targeting the same pMHC occupied consolidated areas of the MDS plot (**Fig. 4e**) and often formed clusters, *e.g.*, clones A3-7/-14/-45, NY-19/-22/-23/-29/-42, and MR-12/-81/-97, suggesting correlations between functional specificity and shared sequence patterns. Intriguingly, clones exhibiting suboptimal functionalities were typically found separate from the cluster centres or at peripheral locations, including TCRm CAR clones with low activations (small disk diameters, such as A3-10, NY-3, NY-25, NY-32, MR-10) and significant cross-reactivities (dash circled, such as NY-2, NY-38, NY-55, NY-58). These spatially discrete clones on the MDS plot were thus excluded from consideration in further studies.

### High specificity and potency of MAGE-A3/A2 and NY-ESO-1/A2 TCRm CARs

A2-restricted MAGE-A3_112-120_ specific TCRm CAR clones A3-45, A3-46 and A3-57 were chosen for in-depth characterizations due to their promising features shown in the initial screening (**Fig. 4b**), including clone abundance, high levels of T activation, and low cross-reactivity. To assess their potencies, binding avidities were measured by flow cytometry with MAGE-A3_112-120_/HLA-A*02:01 pMHC-AF647 tetramer. Results indicated that A3-57 displayed the highest binding avidity to the target pMHC tetramer with an EC_50_ of 60 pM, followed by A3-45 (EC_50_ = 252 pM) and A3-46 (EC_50_ = 598 pM) (**Fig. 5a**). Importantly, their recognitions to the pMHC were peptide-dependent as bindings were fully abolished towards the empty HLA-A*02:01 tetramer. Functional avidities were assessed by NFAT-luciferase activity measurements following APC co-culture pulsed with increasing concentrations of the cognate MAGE-A3_112-120_ peptide. Results showed that all three tested A3 TCRm CAR clones responded to the APC stimulations in a peptide dosage dependent manner, with EC_50_ values of 584, 193 and 540 nM for A3-45, A3-46, and A3-57 respectively (**Fig. 5b**). Notably, A3-46 exhibited the lowest binding avidity but highest functionality among these clones. Similar experiments with NY-ESO-1/A2 specific TCRm CAR clones NY-9, NY-17 and NY-24 revealed their binding avidities of 33, 21, and 34 pM towards NY-ESO-1_157-165_/HLA-A*02:01 tetramer (**Fig. 6a**), and functional avidities of 50, 80, and 37 nM upon stimulation with NY-ESO-1_157-165_ peptide pulsed APCs (**Fig. 6b**). Collectively, these avidity measurements suggested that isolated TCRm CARs exhibited potent and specific recognitions to the cognate peptide loaded HLA with functional avidities ranging from ∼40 to ∼600 nM, comparable to that of typical TCRs in the presence of CD8 coreceptor.

**Figure 5.**
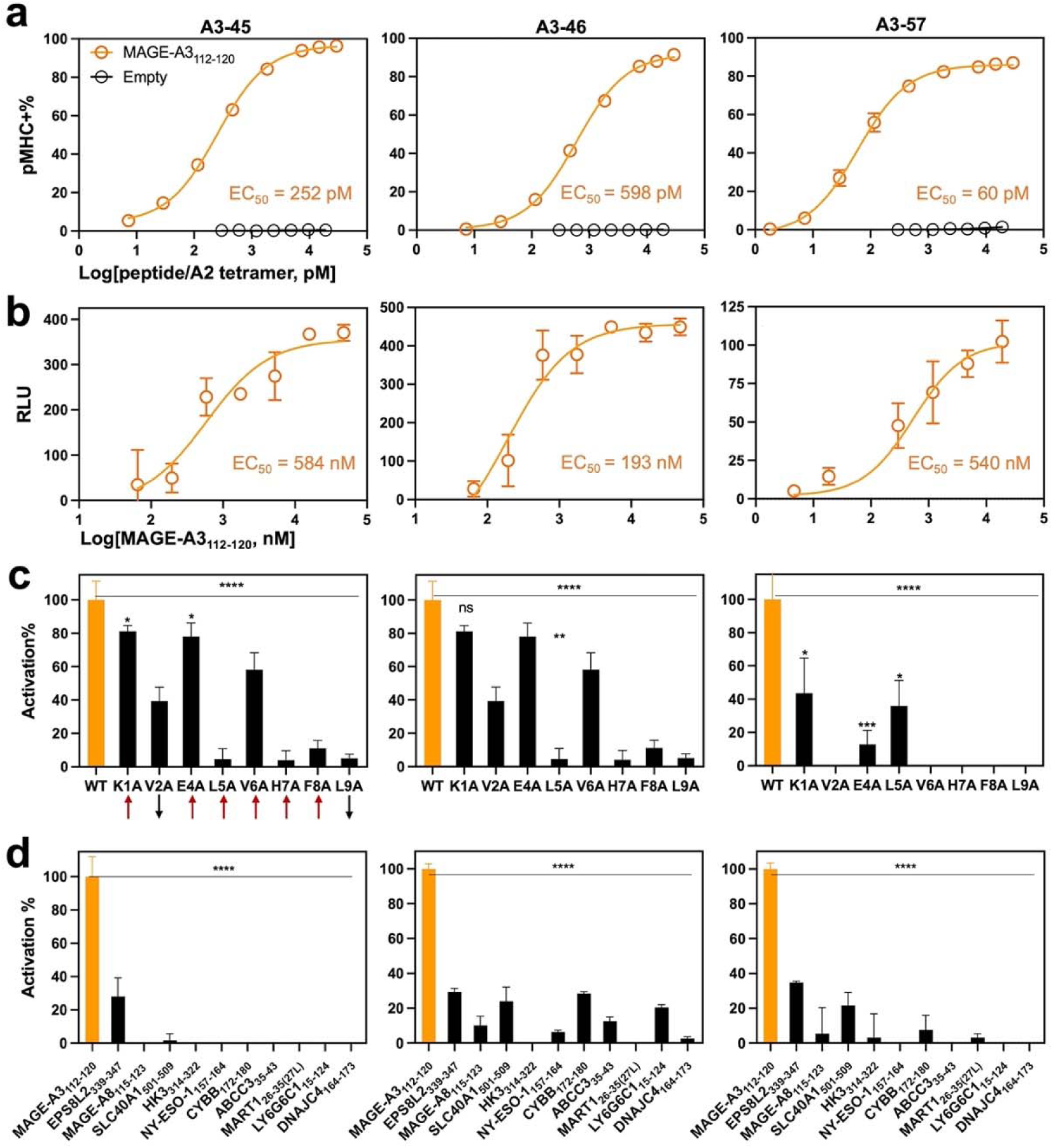
Potency and specificity of MAGE-A3/A2 TCRm CARs. (A3-45, left; A3-46, middle; and A3-57, right columns). **a**, Binding avidity. CAR transduced Jurkat cells were stained with serially diluted MAGE-A3_112-120_/HLA-A*02:01 pMHC-AF647 tetramer and percentages of pMHC^+^ cells were assessed by flow cytometry (n=2). **b**, Functional avidity. CAR transduced Jurkat NFAT-luciferase cells were overnight cocultured with 293F cells pulsed with serially diluted MAGE-A3_112-120_ peptide and the generated bioluminescence signals were measured (n=3). **c**, Functional specificity against alanine mutated peptides (n=3). Upward-, downward-and sideways-facing amino acids are indicated by arrows (PDB=3V5D). **d**, Functional specificity against cross-reactive and irrelevant peptides (n=2). Potential cross-reactive peptides were predicated by sCRAP (**Fig. S8**) [Yarmarkovich *et al*., 2023]. Peptide affinities to HLA-A*02:01 were predicted by NetMHC4.0 (**Fig. S8**) [Andreatta & Nielsen, 2016]. In **c-d**, 293F cells were pulsed with 2.5 µM of tested peptides. Statistical analysis was conducted with one-way ANOVA with Dunnet’s test. ns, not significant; *, p≤ 0.0332; **, p≤ 0.0021; ***, p ≤ 0.0002; ****, p≤ 0.0001.

**Figure 6.**
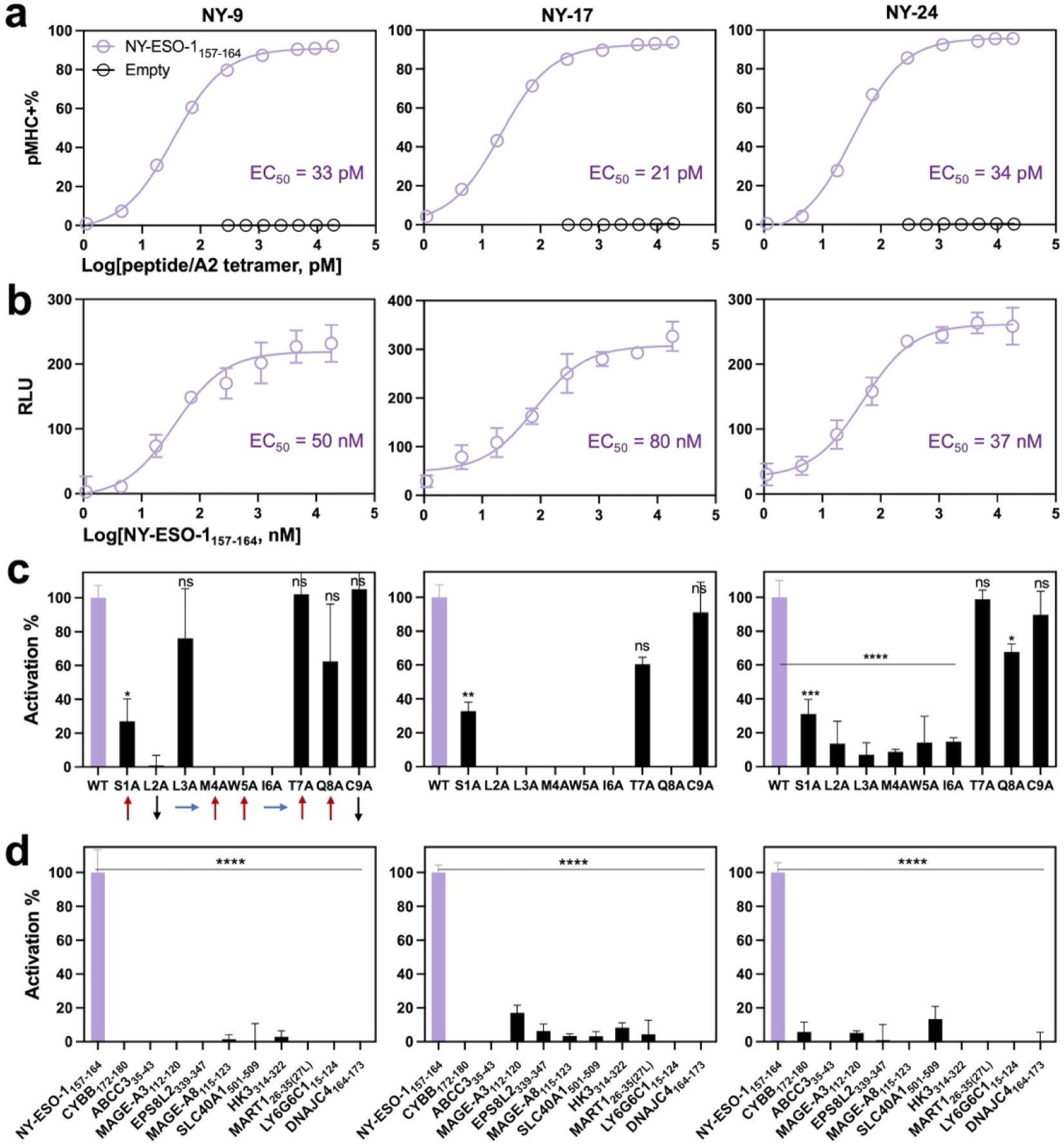
Potency and specificity of NY-ESO-1/A2 TCRm CARs. (NY-9, left; NY-17, middle; and NY-24, right columns). **a**, Binding avidity. CAR transduced Jurkat cells were stained with serially diluted NY-ESO-1_157-165_/HLA-A*02:01 pMHC-APC tetramer and percentages of pMHC^+^ cells were assessed by flow cytometry (n=2). **b**, Functional avidity. CAR transduced Jurkat NFAT-luciferase cells were overnight cocultured with 293F cells pulsed with serially diluted NY-ESO-1_157-165_ peptide and the generated bioluminescence signals were measured (n=3). **c**, Functional specificity against alanine mutated peptides (n=2). Upward-, downward-and sideways-facing amino acids are indicated by arrows (PBD=1S9W) [Webb *et al*., 2004]. **d**, Functional specificity against cross-reactive and irrelevant peptides (n=2). Potential cross-reactive peptides were predicated by sCRAP (**Fig. S8**) [Yarmarkovich *et al*., 2023]. Peptide affinities to HLA-A*02:01 were predicted by NetMHC4.0 (**Fig. S8**) [Andreatta & Nielsen, 2016]. In **c-d**, 293F cells were pulsed with 2.5 µM of tested peptides. Statistical analysis was conducted with one-way ANOVA with Dunnet’s test. ns, not significant; *, p≤ 0.0332; **, p≤ 0.0021; ***, p ≤ 0.0002; ****, p≤ 0.0001.

As cross-reactivity can lead to severe adverse events in pMHC targeted T cell therapies [Linette *et al*., 2013; Morgan *et al*., 2013], it is important to examine functional selectivity of isolated TCRm CARs. We first applied alanine scanning on the target peptide to study amino acid position sensitivity. APCs loaded with alanine substituted peptides were used to assess activation of CAR transduced Jurkat NFAT-luciferase cells by measuring secreted luciferase activity after coculture. For all three tested A3-CAR clones, single substitutions at any non-anchor positions of MAGE-A3_112-120_ peptide resulted in significant reduction of T cell activation. Taking A3-57 as example, alanine mutation of E4A, V6A, H7A or F8A fully abolished the recognition (**Fig. 5c**), suggesting its plausible epitope. For the three NY-CAR clones, alanine scan on NY-ESO-1_157-164_ peptide revealed that the key residues protruding from the HLA cleft [Webb *et al*., 2004], including S1, M4, W5, I6, and Q8, were substantially involved in binding with these CARs (**Fig. 6c**). Overall, the isolated TCRm CARs interacted with 4-6 out of 7 non-anchor residues, highlighting their selectivity at par or higher than typical 3 or 4 residues that interact with TCRs.

Next, potential cross-reactive antigen presentations on HLA-A*02:01 were predicted for each peptide target by sCRAP [Yarmarkovich *et al*., 2023], which considered sequence similarity, gene expression in normal tissues, and pMHC binding affinity. For MAGE-A3_112-120_, four human peptides derived from EPS8L2, MAGE-A8, SLC40A1, and HK3 were identified (**Fig. S8**), of which EPS8L2_339-347_ and MAGE-A8_115-123_ had binding affinities lower than that of the cognate MAGE-A3_112-120_ as predicted by NetMHCpan [Andreatta & Nielsen, 2016]. Additional 6 peptides, including NY-ESO-1_157-164_, MART-1_26-35_ and their potential cross-reactive predictions, were also included in the Jurkat NFAT-luciferase activation assays to assess the cross-reactivity. Among the 10 non-cognate peptides tested, A3-45 CAR only weakly responded to EPS8L2_339-347_ at 28% activation compared to the cognate antigen peptide at the same condition (**Fig. 5d**). Similarly, negligible or dramatically reduced activations were observed for A3-46 and A3-57 CARs towards all the non-cognate peptides tested. For NY-ESO-1_157-164_, potential cross-reactive peptides CYBB_172-180_ and ABCC_35-43_ were applied (**Fig. S8**), and results showed that none of NY TCRm CARs reacted with any non-cognate peptides tested (**Fig. 6d**). Overall, alanine scanning and studies with predicted cross-reactive peptides collectively suggested that isolated TCRm CARs mediated highly specific T cell activation towards their cognate pMHC.

### *In vitro* cytotoxicity of MAGE-A3/A2 and NY-ESO-1/A2 TCRm CARs

Primary T cells were retrovirally transduced with the identified MAGE-A3/A2 specific TCRm CARs and co-cultured with human melanoma A-375 or osteosarcoma SaOS-2 cells, both HLA-A2^+^ and MAGE-A3^+^ double positive. To examine the CAR-T cell response, we measured the releases of a key cytotoxic cytokine interferon-γ (IFN-γ). In comparison to non-transduced T cells, A3-45, A3-46, and A3-57 TCRm CAR-T cells secreted high amounts of IFN-γ (up to 1250 pg/ml) upon cocultures with the target cells, indicating CAR mediated T cell activation (**Fig. 7a**). Their functional specificity towards MAGE-A3/HLA-A*02:01 was evident from the observation that only basal levels of IFN-γ were induced by HLA-A2^-^ / MAGE-A3^-^ A-549 or HLA-A2^+^ / MAGE-A3^-^ 293T cells (**Fig. 7a**). Cytotoxic effector functions of A3-45, A3-46, and A3-57 TCRm CAR-T cells were assessed by measuring the enzymatic activities of intracellular proteases released from membrane-compromised target cells. At the effector to target cell ratio (E:T) of 10:1, 27-43% targeted tumor cells were lysed, representing 6-to 10-fold increases over the control cell lines (**Fig. 7b**). Such HLA restricted, cognate peptide specific T cell activation and cytotoxicity were equally achieved with NY-9, NY-17 and NY-24 TCRm CAR-T cells – they exhibited potent and selective IFN-γ secretion and cytotoxicity against HLA-A2^+^ and NY-ESO-1^+^ double positive A-375 and SaOS-2 cells, whereas their effects on A-549 (HLA-A2^-^ / NY-ESO-1^-^) and 293T (HLA-A2^+^ / NY-ESO-1^-^) were minimal (**Fig. 8a-b**). With elevated E:T ratios, the cytotoxicity of TCRm CAR-T cells significantly enhanced as expected (**Fig. 7c**, **Fig. 8c**), *e.g.*, specific lyses rising from 24% to 41% and 54% when the NY-24 CAR-T to SaOS-2 cell ratio increasing from 5 to 10 and to 20:1. The cytotoxic functions of TCRm CAR-T cells were further confirmed with primary T cells from different donors and visualized with live-cell imaging, *e.g.*, A3-45 and A3-46 TCRm CAR-T cells clustered around A-375 cells and induced tumor cell lysis (**Fig. S9**). Collectively, these results indicated the efficacy and specificity of TCRm CAR-T in targeting MAGE-A3/HLA-A2 or NY-ESO-1/HLA-A2 positive tumor cells, suggesting their therapeutic potential in cancer treatment.

**Figure 7.**
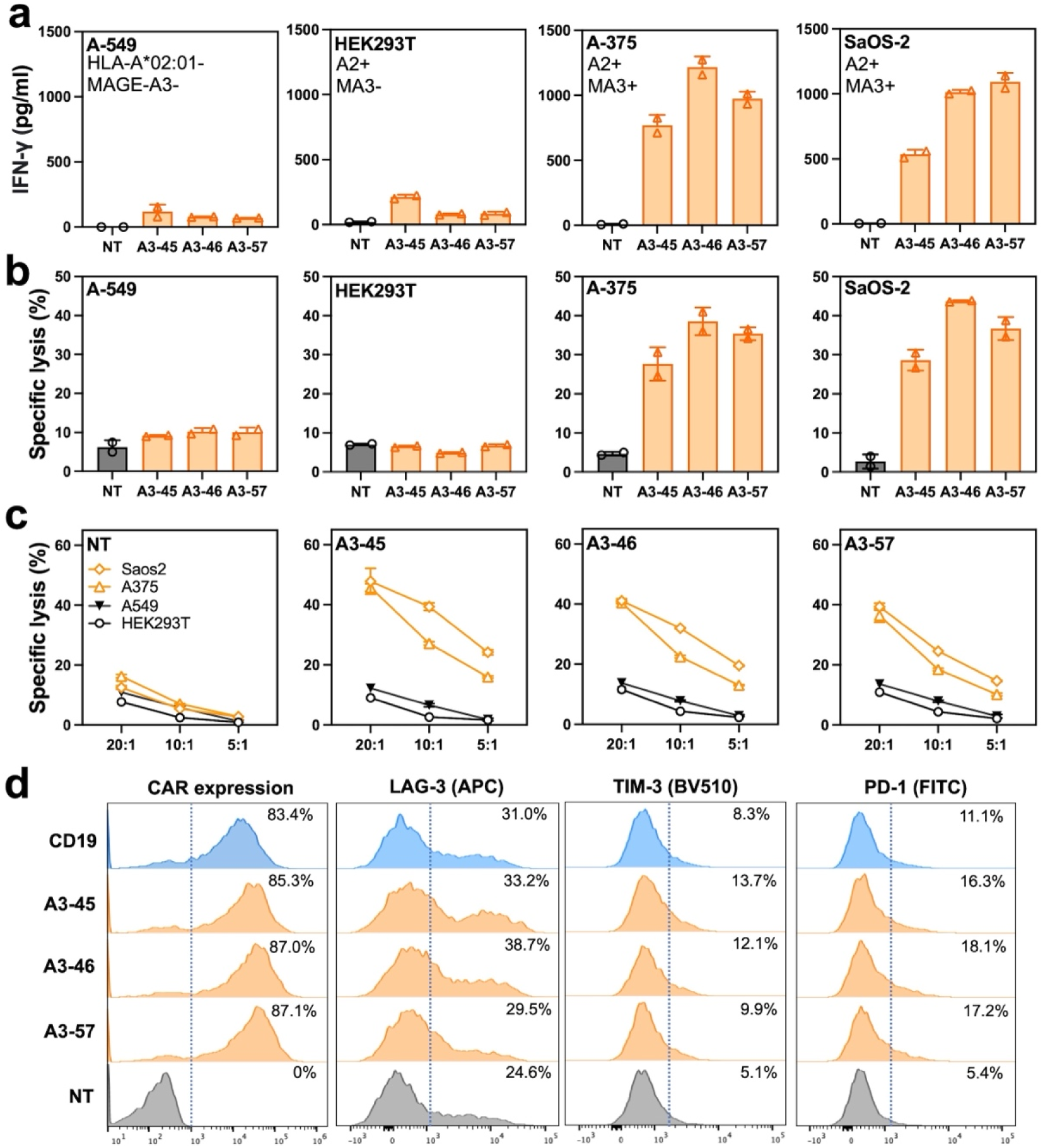
Specific, potent and persistent primary T cell activity of MAGE-A3/A2 TCRm CARs. **a**, IFN-γ release measured by ELISA upon cocultures with target cell lines at an E:T of 5:1. (n=2). **b**, Cytotoxic activity of A3 TCRm CAR-T cells against the indicated tumor cell lines assessed by intracellular protease activity assays at an E:T of 10:1. Tested cells included A-549 (HLA-A2^-^, MAGE-A3^-^), HEK293T (HLA-A2^+^, MAGE-A3^-^), A-375 (HLA-A2^+^, MAGE-A3^+^), and SaOS-2 (HLA-A2^+^, MAGE-A3^+^). Non-transduced T cells were applied as controls. (n=2). **c**, Cytotoxic activity at different E:T ratios. **d**, Flow cytometry histograms showing expression of CAR, LAG-3, TIM-3, and PD-1 on day 10 post transduction. Non-transduced T cells and CD19 CAR-T cells with low tonic signaling were tested as well.

**Figure 8.**
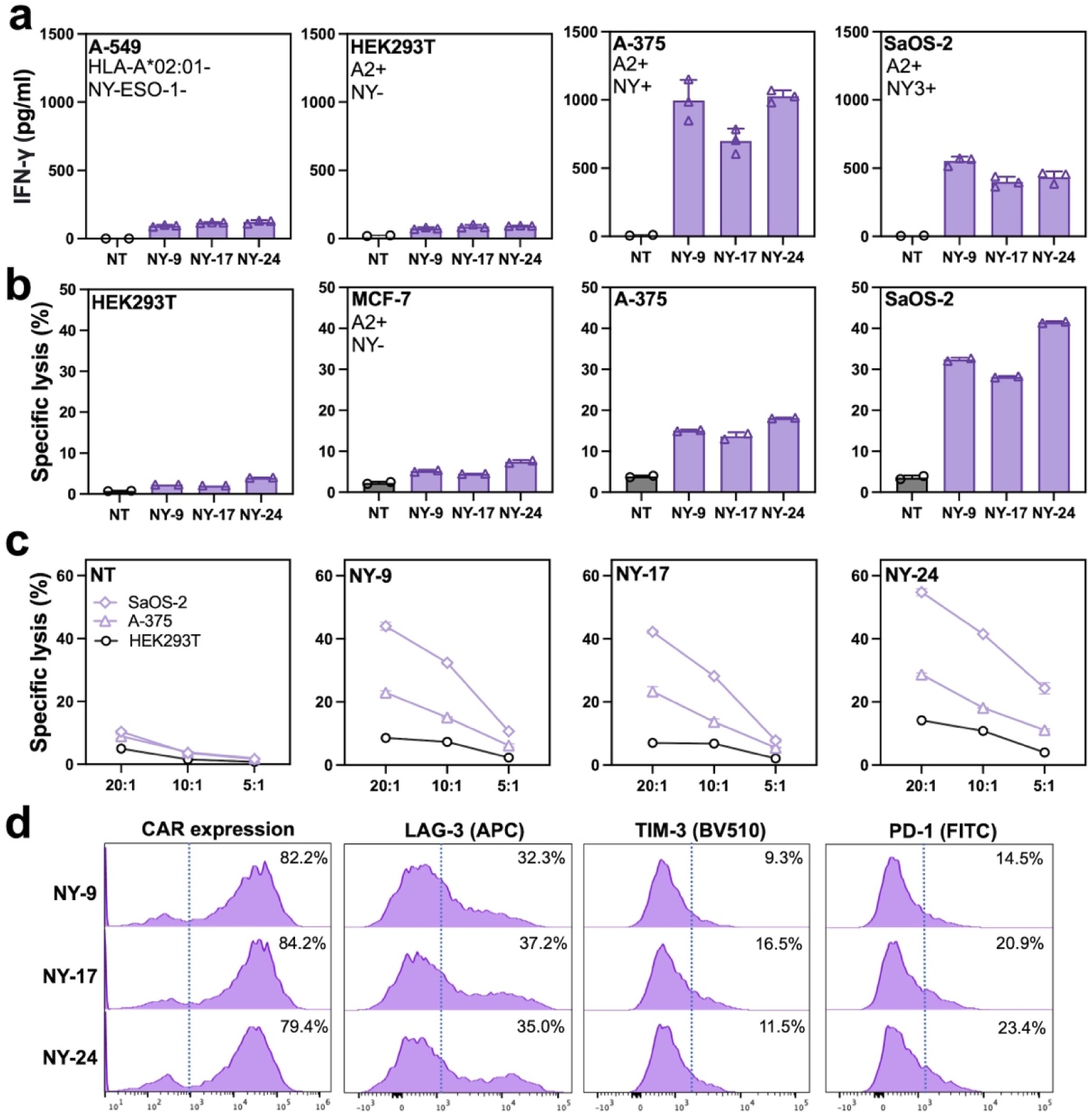
Specific, potent and persistent primary T cell activity of NY-ESO-1/A2 TCRm CARs. **a**, IFN-γ release measured by ELISA upon cocultures with target cell lines at an E:T of 5:1. (n=3). **b**, Cytotoxic activity of NY TCRm CAR-T cells against the indicated tumor cell lines assessed by intracellular protease activity assays at an E:T of 10:1. Tested cells included A-549 (HLA-A2^-^, NY-ESO-1^-^), HEK293T (HLA-A2^+^, NY-ESO-1^-^), MCF-7 (HLA-A2^+^, NY-ESO-1^-^), A-375 (HLA-A2^+^, NY-ESO-1^+^), and SaOS-2 (HLA-A2^+^, NY-ESO-1^+^). Non-transduced T cells were applied as controls. (n=2). **c**, Cytotoxic activity at different E:T ratios. **d**, Flow cytometry histograms showing expression of CAR, LAG-3, TIM-3, and PD-1 on day 10 post transduction. Non-transduced T cells and CD19 CAR-T cells with low tonic signaling were shown in Fig. 7d.

Strong tonic signaling causes rapid T cell exhaustion and thus impairs persistence of antitumor cytotoxicity [Long *et al*., 2015]. On the other hand, inefficient tonic signaling can also lead to reduced T cell effector function [Singh *et al*., 2021]. Therefore, a balanced cell-autonomous response is desired for an efficacious CAR-T therapy. To assess the basal state of isolated A3 and NY TCRm CARs, transduced primary T cells were cultured in the absence of tumor antigen stimulation and the key exhaustion markers were monitored for comparison with those of CD19 CAR, which has been proven highly efficacious with low tonic signaling [Tousley *et al*., 2023; Chen J *et al*., 2023]. On day 10 post-transduction, 79-87% of cells displayed TCRm CARs, consistent to that of CD19 CAR-T at 83% (**Fig. 7d**, **Fig. 8d**). We found that for the six A3 and NY TCRm CAR clones, the expression of lymphocyte activation gene 3 (LAG-3) ranged from 29% to 38%, on par with 31% for CD19 CAR (**Fig. 7d**, **Fig. 8d**). Similarly, expressions of T cell immunoglobulin and mucin domain-containing protein 3 (TIM-3) and programmed cell death protein 1 (PD-1) were measured at 12.2±2.6% and 18.4±3.2% among TCRm CAR-T cells, comparable to CD19 CAR-T at 8.3% and 11.1% respectively (**Fig. 7d**, **Fig. 8d**). Non-transduced T cells were also examined, and the results confirmed the above background but low tonic signaling for both CD19 and all tested TCRm CARs. On day 20, expressions of these three inhibitory receptors increased across transduced TCRm and CD19 CAR-T cells as expected. However, no significant differences were observed by comparing TCRm CARs with CD19 CAR (**Fig. S10**). In addition, the levels of TCRm CAR display were largely maintained as well. Overall, analysis of three exhaustion markers collectively indicated the persistence of TCRm CARs without signs of early T cell exhaustion, presumably attributing to the efficient removal of clones showing spontaneous activation during our functional TCRm CAR screening (**Fig. 3**).

## DISCUSSION

CAR-T therapy has received remarkable clinical successes in the treatment of hematological malignancies, but its impact on solid tumor remains modest, largely due to the reliance on cell surface antigens [Baulu *et al*., 2023]. Given that most tumor-specific antigens reside in cytoplasm or nucleus and thus are only accessible to immune system through the presentation of peptide-MHC complexes, the development of adoptive cell therapies targeting pMHC of intracellular cancer antigens provides promising opportunities to treat solid tumors. Here, we describe the establishment and applications of a platform technology on efficient discovery of TCR mimic CARs that not only specifically bind tumor antigen pMHC complexes but also effectively elicit potent and persistent T cell activation. Panels of highly functional CAR clones with minimal cross-reactivity and low tonic signaling were rapidly identified from human naïve antibody repertories towards intracellular tumor-associated antigens including MAGE-A3, NY-ESO-1, and MART-1. Measurements of functional avidity, cytokine release, and in vitro cytotoxicity collectively validated the efficacy and specificity of TCRm CAR-T cells, suggesting their therapeutic potential against solid tumors. Intriguingly, CDR distance analysis of the isolated TCRm CAR clones revealed correlations between functional specificity and conserved sequence patterns, a characteristic observed in TCRs [Dash *et al*., 2017]. This finding could set the foundation to generate training data for machine learning model on CAR-pMHC specificity prediction. In addition, peptide-dependent avidities and alanine scanning results jointly suggested that the epitopes of TCRm CARs, like TCRs, involved multiple key non-anchoring residues especially the ones protruding from the HLA cleft. Despite these resemblances to TCRs, further biochemistry and structure biology studies are warranted to fully elucidate the mechanism of TCRm CARs on pMHC recognition. Nonetheless, TCRm CAR-T offer several advantages over TCR-T such as no liability to TCR mismatch, manufacture process standardization, and versatility for the development of T cell engagers.

One key component of our TCRm CAR discovery approach is recombinase-mediated high-efficiency transgene integration in immortalized T cells. Using large serine recombinase Pa01 [Durrent *et al*., 2023], up to 40% integration efficiencies of multi-kilobase DNA payloads were achieved in Jurkat cells for the first time with desired features of monoclonality, site-specificity, fidelity, and high cell viability. This development presents a clear advancement over traditional methods of T cell genetic modification – *e.g.*, viral vectors are limited by issues of biases introduced during packaging, random integration, multiple transgenes per cell, and variation on transgene expression; and CRISPR-based technologies are often associated with low integration efficiency and undesigned mutations. In contrast, our Pa01-mediated integration occurs at specific genomic locus – exon 11 of NGDN on chromosome 14 in the E11 cell line and intron 3 of EIF3H on chromosome 8 in the B4 cell line (**Fig. S3**). Such site-specific monoclonal integration of a library of transgenes ensures uniform transcription level, particularly beneficial for combinatorial library screening. With integration efficiency of ∼25% for CAR transgenes and by electroporation of 8.0×10^7^ Pa01-Jurkat cells, monoclonal libraries of 2.0×10^7^ CAR clones were constructed in immortalized T cells. This library size represents two orders of magnitude larger than the T cell library diversities reported so far, allowing *de novo* discovery of functional TCRm CARs from a human naïve antibody library. Moreover, as the electroporation approach is largely scalable, construction of libraries exceeding 10^8^ diversity is feasible by processing 10^9^ cells. Looking forward, several strategies could further enhance the efficiency and robusticity of recombinase-mediated transgene integration, such as reducing the donor plasmid size with minicircle DNA, incorporating the landing pad into a genomic safe harbor, and applying cell cycle synchronization [Chen C *et al*., 2023].

Another crucial aspect of our system is the employment of T cell activation-based screenings. Our data clearly evidence that the combination of negative and positive screenings is both essential and sufficient for the complete removal of cross-reactive or self-activated clones and the efficient isolation of functional TCRm CARs specific to cognate pMHC. Clone characterizations indicated that over the three tested TAA targets, in average 89% of isolated unique CARs were able to trigger pMHC-restricted T cell activation, an astounding hit rate unfeasible for approaches solely based on pMHC binding such as phage panning or yeast display. Furthermore, the six in-depth analyzed TCRm CARs all exhibited potent functionality (avidity EC_50_ of 40-600 nM) and efficacious and specific cytokine release and cytotoxicity. In addition, these TCRm CARs showed low tonic signaling and negligible cross-reactivities towards 10 non-cognate peptides including the top candidates predicted by sCRAP [Yarmarkovich *et al*., 2023]. All these favorable characteristics suggest potent, highly specific, and persistent T cell functions, which will be further validated by *in vivo* efficacy and safety studies. Taking advantage of this *in vitro* functional selection approach, we expect an extended application where single peptide antigen displayed by multiple HLA allotypes can be collectively used for the generation of peptide-centric CARs across HLAs [Yarmarkovich *et al*., 2023].

In summary, this study combines large-diversity monoclonal library construction in T cells with functional positive / negative screenings, and demonstrates rapid discovery and validation of TCRm CARs specific to HLA-A*02 restricted tumor-associate antigens. This platform technology is readily applicable for the generation of TCRm CARs toward neoantigens and peptide antigens presented by HLA allotypes other than A2. Beyond oncology, this development should have applications in infectious diseases, because eliciting cytotoxic T cells specific to the pathogenic pMHC is critical for pathogenesis suppression and effective virus elimination [Varela-Rohena *et al*., 2008]. Furthermore, targeting autologous pMHC has shown the promise to delay and even prevent the progression of autoimmune diseases such as type 1 diabetes (T1D) [Zhang *et al*., 2019; Uenishi *et al*., 2024]. In addition to CAR, construction of large-diversity monoclonal libraries in immortalized T cells can be extended for the studies of functional TCRs (**Fig. 1e**, **Fig. 2b-c**), T cell antigens (such as SABR) [Joglekar *et al*., 2019; Zdinak *et al*. 2024], and orthogonal cytokine/receptor pairs [Sockolosky *et al*., 2018; Li & Lim, 2020]. Overall, we envision that the technology described here has the potential to substantially advance the development of diverse immunotherapy modalities including CAR-T, TCR-T, and T cell engagers targeting HLA restricted antigen peptides for the treatment of solid tumors, infections, and autoimmune diseases.

## METHODS

### Molecular cloning

The 1G4 TCR encoding gene was amplified from pTR262 (Addgene, 112022) and cloned into the donor plasmid Pa01attP-mCherry (Addgene, 193466) with BsrGI and XhoI, leading to the donor plasmid Pa01attP-1G4 TCR. The scFv genes of 3M4E5 and MA2 were synthesized as gBlock (IDT). The 28z gene construct including CD8 hinge, CD28 transmembrane domain, CD28 costimulatory and CD3z signaling domains, was amplified from pSLCAR-CD19-28z (Addgene, 135991) and linked to the downstream of 3M4E5 scFv via overlap PCR. The truncated LNGFR gene (ΔLNGFR, LNGFR excluding intracellular region) was amplified from pPRIME-CMV-LNGFR-FF3 (Addgene, 11666). 3M4E5 CAR fused with ΔLNGFR via a F2A self-cleaving peptide was cloned into Pa01attP-1G4 TCR with BsrGI and XhoI, named as Pa01attP-3M4E5 CAR-F2A-LNGFR. The MA2 CAR donor plasmid was obtained through site directed mutagenesis of Pa01attP-3M4E5 CAR-F2A-LNGFR. The CD19 CAR lentiviral expression plasmid pSLCAR-CD19-28z was modified to remove the CD19 scFv and replace the EGFP with mCherry gene, resulting in pSL-28z-mCherry. The SFG retroviral plasmid, envelope plasmid RD114 and packaging plasmid Gag-pol pEQ-Pam3 were generously gifted by Dr. Norihiro Wantabe of Baylor College of Medicine. The signaling peptide and 28z-mCherry construct were cloned into SFG retroviral plasmid cut with XhoI and SphI, resulting in SFG-28z-mCherry. The scFv gene was cloned into SFG-28z-mCherry with NgoMIV and NotI leading to SFG-CAR-2A-mCherry. All the restriction enzymes, polymerases and DNA ligase were purchased from NEB. All cloned plasmids were confirmed by sanger sequencing.

### Cell culture

Jurkat E6-1 T cell line (ATCC, TIB 152), Jurkat-Pa01 (made in house), and Jurkat-NFAT-EGFP (made in house) cells were cultured in R10 media composed of RPMI1640 medium (Corning, 10-040-CV), supplemented with 10% fetal bovine serum (FBS, Gibco, 10437028), 50 U/ml penicillin, and 50 µg/ml streptomycin (Gibco, 15140122). Jurkat-Lucia NFAT cells (Invivogen, jktl-nfat) were cultured in IMDM media (Cytiva, SH30228.02), supplemented with 10% FBS, 50 U/ml Pen-Strep, and selective antibiotic zeocin was used every alternative passage. A-375 (MD Anderson Cytogenetics and Cell Authentication Core), MCF-7 (ADCC, HTB-22), LentiX-293T cells (Takara, 632180) and HEK293T (ATCC, CRL-3216), A-549 (ATCC, CCL-185), MDA-MB-231 (ATCC, HTB-26) were cultured in DMEM medium (Gibco, 10569010), supplemented with 10% FBS, 50 U/ml Pen-Strep. SaOS-2 (ATCC, HTB-85) cells were cultured in McCoy5A media (ATCC, 30-2007), supplemented with 15% FBS. All cells mentioned above were cultured at 37 °C, in 5% CO_2_ humidified atmosphere. Expi293F (Gibco, A14527) was cultured in Expi293 medium (Gibco, A1435101) supplemented with 40 U/ml penicillin and 40 µg/ml streptomycin at 37 °C, in 8% CO_2_ humidified atmosphere with a shaking speed 130 rpm. Cell numbers and viabilities were measured with 0.4% trypan blue staining followed by detection on Countess 3 automated cell counter (Invitrogen).

### Peptides, peptide MHC complexes and peptide pulse

Peptides NY-ESO-1_157-165_ (SLLMWITQV), MAGE-A3_112-120_ (KVAELVHFL), MART-1(A2L)_26-35_ (ELAGIGILTV), CYBB_172-180_ (TLLAGITGV), ABCC_35-43_ (SLLAWVPCI), EPS8L2_339-347_ (SAAELVHFL), SLC40A1_501-509_ (YLLDLLHFI), HK3_314-322_ (YLGELVRLV), MAGE-A8_115-123_ (KVAELVRFL), LY6G6C_115-124_ (SLAGLGLWLL), DNAJC4_164-173_ (MLAGMGLHYI), and alanine-substituted NY-ESO-1_157-165_ and MAGE-A3_112-120_ were synthesized (GenScript) and dissolved in DMSO at 10 mg/ml. Alexa Fluor 647 or APC conjugated peptide MHC (pMHC-AF647, pMHC-APC) tetramers were obtained from the NIH tetramer core facility. Biotinylated human HLA-A*02:01 (A2) monomer (HLM-H82W3) and MAGE-A3_112-120_/A2 monomer (HLM-H82E3) were purchased from AcroBiosystems. Biotinylated NY-ESO-1_157-165_/A2 and MART1(A2L)_26-35_/A2 pMHC monomers were obtained by loading the corresponding peptide to biotinylated empty A2 monomer following manufacturer’s protocol. Specifically, 20 µl of 1 mg/ml peptides (10 times diluted in PBS) was slowly added to 100 µl of 500 µg/ml biotinylated MHC monomer. The next day, the peptide loaded MHC monomer was purified by ultrafiltration. Tetramers were prepared by adding streptavidin-APC (Agilent Technologies, PJ27S) to the peptide-A2 monomer in dark. For peptide pulsing, Expi293F cells were harvested and resuspended in fresh Expi293 media to a density of 1×10^6^/ml. Peptides were then added to a final concentration of 10 µg/ml. Cells were cultured with shaking for 3 h, washed once with Expi293 media, and ready for co-culture experiments.

### Lentivirus production

4×10^6^ of LentiX-293T cells were seeded in a 10 cm plate. Next day, the media was replaced with 13.5 ml of fresh media. 9.2 µg plasmid of interest, 8.3 µg psPAX and 2.5 µg pMD2.G were mixed with 80 µg PEI MAX 40K (Polysciences, 24765–1) in 1.5 ml of Expi293 media. After resting at room temperature for 15 min, the DNA/PEI mixture was added to the plate dropwise slowly. 12 hours after the transfection, the supernatant was replaced with 15 ml of fresh media. After 48 h, the supernatant was harvest and centrifuged at 1000 g for 10 min. Optionally, the supernatant was concentrated with a 30 kDa protein concentrator (Thermo Scientific, 88531). For packaging in a smaller scale, the DNA amount and cell number were scaled down according to the plate surface area.

### Generation of Jurkat-Pa01 cell lines

The Pa01 landing pad plasmid (Addgene, 184948) was packaged to lentivirus as described above. Two milliliters of 1×10^6^ Jurkat cells were mixed with different doses of lentiviruses in 8 µg/ml polybrene and infected overnight without spinfection. Next day, the cells were cultured in fresh media for 2 more days, and the percentage of EGFP+ cells was detected by flow cytometry. The cells with below 5% EGFP+ population were further expanded, and one week later, EGFP+ cells were sorted out and named as Jurkat-Pa01. To obtain monoclonal cell line, Jurkat-Pa01 polyclonal cells were seeded into 96-well plates at a density of 0.5 cell/well. After 3 weeks of culture, single cell clones were identified and expanded for further study.

### Genotyping of the landing pad genomic locus

Inverse PCR was employed to determine the landing pad copy number per cell and its genomic locus. Briefly, 30 ng genomic DNA was digested with each of the restriction enzymes (TaqI, EagI-HF, PciI, MfeI-HF, and NcoI-HF) in a 10 μl reaction for 4 h followed by heat inactivation at 80°C for 20 min. The digestion mixture was then added with 40 μl of T4 DNA ligase buffer containing 64 units of T4 DNA ligase and incubated at 16°C overnight. The self-ligated products were amplified using two rounds of PCR. The nested PCR products were examined through gel electrophoresis followed by gel extraction and Sanger sequencing.

### Electroporation

Electroporation was performed on MaxCyte STx following the manufacture’s protocol. Briefly, Jurkat cells were washed twice with and resuspended in Electroporation Buffer (MaxCyte) at a density of 1×10^8^/ml. 2× 10^6^ cells were mixed with 4 µg of donor plasmid and electroporated using the OC-25×3 processing assembly (PA). Cells were recovered at 37 °C for 30 min then added with fresh media for cultivation at a density of 1×10^6^/ml.

### Flow cytometry analysis

0.5 µg/ml anti-NGFR-PE (clone ME20.4, BioLegend, 345105), 10 µg/ml pMHC-AF647 tetramers and 0.5 µg/ml anti-CD69-PE (clone FN50, BioLegend, 985202) were generally used for staining. All stainings were performed in 0.5% BSA in PBS buffer (PBSA) for 30 min at 4 °C in dark and then washed with PBSA three times before flow cytometry analysis in a Beckman CytoFlex SRT. In tetramer binding assays, 10^5^ Jurkat CAR-T cells were stained with MAGE-A3_112-120_/A2 pMHC-AF647 or NY-ESO-1_157-165_/A2 pMHC-APC at 1 pM-30 nM.

### scFv biopanning

A large naïve phage display scFv library of ∼10^11^ diversity [Ku *et al*., 2021] was subjected to two round of biopanning towards specific pMHC. In all depletion or selection steps, phages were incubated with biotinylated pMHC monomer for 1 h followed by incubation with magnetic beads (MyOne Streptavidin T1 Dynabeads, Invitrogen, pre-blocked with 3% BSA in PBS) for an additional hour. In the first round, a total of 10^13^ copies of phages were depleted against 5 µg of empty MHC monomer followed by selection with 5 µg target pMHC monomer. The phage bound beads were washed with PBS containing 0.05% Tween-20 (PBST) 5 times followed by 2 PBS washes. In the second round, a total of 10^12^ copies of phages were sequentially depleted against empty T1 beads, 5 µg of empty MHC monomer, mixture of two irrelevant pMHC monomers (2.5 µg each) prior to the selection for binding with 2.5 µg of the target pMHC monomer. The phage bound beads were washed 8 times with PBST and twice with PBS. In each round, the remaining bound phage were eluted by 10-min incubation with 200 µl fresh 100 mM TEA followed by neutralization with 100 µl 1 M Tris-HCl (pH 7.5). The eluted phages were recovered by 10 ml fresh prepared log-phase TG1 and rescued using the M13K07 helper phage prior to culture on 2×YT/Amp agar plate.

### TCRm-CAR library construction in Jurkat-Pa01 and selection

The scFv gene pool was amplified by 6-cycle of PCR from round 2 output phagemids followed by cloning into the donor plasmid Pa01attP-CAR-F2A-LNGFR with NEBuilder HiFi DNA assembly. 8 × 10^7^ Jurkat-Pa01 B4 cells were electroporated with 160 µg TCRm-CAR library donor plasmid in OC-400 RUO. On day 7 post electroporation, library cells were harvested and incubated with 0.4 µg/ml of biotinylated anti-LNGFR (clone ME20.4, BioLegend, 345122) at 4 °C for 30 min followed by MACS (Miltenyi Biotec) following manufacture’s protocol. The bead bound cells were expanded for FACS as described in previous section with modifications. Specifically, in the binding based selection round, cells were stained with 5 µg/ml target pMHC-AF647 tetramer and 25 ng/ml anti-LNGFR-PE. Both PE+ and AF647+ cells were sorted for expansion. In the activation-based selection, the CAR displayed Jurkat cells were cocultured with equal number of either target or irrelevant peptide-pulsed 293F cells overnight for selection or counter-selection, respectively. The next day, cells were stained with 2.5 µg/ml target pMHC-AF647 tetramer and 0.5 µg/ml anti-CD69-FITC FN50 (Biolegend, 310904) for FACS. In a counter-selection, FITC negative while AF647 positive cells were sorted; in a positive selection, cells with high FITC and AF647 signals were sorted.

### TCRm CAR clone identification and screening in Jurkat NFAT-GFP

With cells collected from the final round of FACS, genomic DNA was extracted with QIAamp DNA Micro Kit (Qiagen, 56304). From the gDNA, scFv genes were amplified and assembled into the pSL-28z-mCherry linearized by BamHI and KpnI. After transformation into NEBstable, single colonies were randomly picked for minipreps and Sanger sequencing. The identified unique TCRm-CAR clones were then packaged to lentivirus in a 96-well plate. For transduction, 180 µl of Jurkat NFAT-GFP in 5×10^5^ cells/ml containing 8 µg/ml polybrene were mixed with 20 µl of the virus in a 96-well plate followed by centrifugation at 800 g 30 °C for 90 min. 2 days after transduction, 90 µl of the transduced Jurkat cells were added with same volume of peptide-pulsed Expi293F cells (1×10^6^ cells/ml) in R10 media and co-cultured overnight for next day flow cytometry analysis.

### CAR distance measure and visualization

The distance between any two CAR clones is defined as accumulated similarity-weighted mismatch distances between their respective CDRs. For each CDR pair, alignment was first performed by using pairwise2 module (Biopython) with the gap opening penalty at -10 and the gap extension penalty at -0.5. From the resulting optimal alignment, mismatch distances for each CDR pair were then calculated using the BLOSUM62 substitution matrix with the following rules of TCRdist [Dash *et al*., 2017]: distance (a,a) = 0; distance (a,-) = 4; distance (a,b) = min (4, 4-BLOSUM62 (a,b)), where 4 is 1 unit greater than the most favorable BLOSUM62 score for a mismatch, a and b are amino acids, and “-” is a gap. For each CDR pair, the total mismatch distance was computed as the sum of individual distance across the alignment. These distances were compiled into an array for each CAR pair. To account for the greater role of the CDR3 regions in antigen recognition, a weight factor of 3 is applied to CDR3s. A distance matrix between all CARs was computed accordingly and used for multidimension scaling to visualize the distances among CARs in a two-dimensional space.

### Jurkat NFAT-luciferase activation assay

2.5×10^5^ Jurkat-Lucia NFAT cells were transduced with 20 µl lentivirus packaged with TCRm CARs. On day 3 after transduction, 2×10^4^ transduced Jurkat cells were cocultured with equal number of Expi293F cells pulsed with 4× serial diluted peptides in a 96-well plate. After overnight culture, the plate was centrifuged at 250 ×g for 3 min. From each well, 20 µl supernatant was transferred to a 96-well optical clear bottom black plate mixed with 50 µl of prepared QUANTI-Luc™ 4 Lucia/Gaussia reagent to detect secreted luciferases per manufacturer’s instruction. For alanine substituted and cross-reactive peptides, 5×10^4^ Jurkat CAR-T cells were coculture with Expi293F cells pulsed with 2.5 µg/ml peptide. All measurements were performed on a BioTek Cytation 5.

### Retroviral production

2×10^6^ of HEK293T cells were seeded in a 10 cm plate. Next day, the cell should be 60-70% confluent. Transfection mixture was prepared in DMEM without any supplements. In a sterile 1.5 ml Eppendorf tube, 30 µl of GeneJuice transfection reagent (Sigma Aldrich, 70967) was added directly to 420 µl of DMEM and mixed by gently tap. In a separate tube, a total of 10 µg DNA was added to 50 µl of DMEM, including 3.75 µg pEQ-Pam3, 2.5 µg pRD114 and 3.75 µg SFG-CAR-2A-mCherry. Dropwise add the GeneJuice/DMEM mixture to the tube containing the DNA. The mixture was incubated for 15 min at room temperature and then added to the plate dropwise slowly. After 12 h, the supernatant was replaced with 10 ml of fresh media. After 48 h, the supernatant was harvest and filtered with 0.45 µm filter. In labeled tubes add 1 ml aliquots of supernatant to store at -80 °C.

### Transduction of human primary T cells with TCRm CARs

A 24-well non-treated tissue culture plate (StemCell, 100-0097) was coated with 1 ml of 1 µg/ml anti-CD3 (OKT3 clone) and CD28 at 4 °C overnight. The next day, 1 million human peripheral blood mononuclear cells (PBMCs, STEMCELL, 70025) were added to the plate and incubated at 37°C, 5% CO_2_ for 2 days. The retroviral plasmid carrying the TCRm CAR expression cassette was packaged to retrovirus as described above. One day prior transduction, a new 24-well non-treated tissue culture plate was coated with 10 µg retronectin (Takara, T100A) in 1 ml PBS per well, sealed with parafilm and kept at 4 °C overnight. Next day, the retronectin solution was removed and 1 ml of retrovirus packaged with TCRm CAR was added per well followed by centrifugation at 4,600 × g for 1 h at 30 °C. The supernatant was then aspirated and 5×10^5^ preactivated human PBMCs were added and cultured in 1 ml R10 media supplemented with 100 U/ml human IL-2. The plate was centrifuged at 800 ×g, 30 °C for 10 min. 48 hours after transduction, the media was replaced with fresh R10 media with IL-2 for continuous expansion of transduced CAR-T cells.

### ELISA

IFN-γ release upon activation of CAR-T cells was measured by ELISA MAX^TM^ standard set (Biolegend, 430101). 2.5×10^4^ target cells were overnight seeded in 96 well plates and cocultured with transduced CAR-T cells at E:T ratios of 2:1, 5:1, or 10:1 for 18 hrs. After centrifugation, 100 µl of media were collected for assays, and the amounts of IFN-γ were determined against standard curves generated with 0-2000 pg IFN-γ. Non-transduced T cells were used as the control.

### Primary T cytotoxicity assay

Cytotoxicity assays were performed using CytoTox-Glo (Promega, G9290) following manufacturer’s instruction. In brief, 2.5×10^4^ target cells were overnight seeded in white 96 well optical bottom tissue culture plates (Thermo Fisher 165306) and cocultured with transduced CAR-T cells or non-transduced T cells at E:T ratios of 5:1, 10:1 or 20:1 for 18 hr. 50 µl of assay reagent was then added to cell cultures, mixed by orbital shaking, and incubated at room temperature for 15 min. Signals of bioluminescence were measured by using a BioTek Cytation 5. Target cells cultured and processed without effector T cells were used to generate background signals, and target cells lysed by digitonin were used to set maximal lysis levels. Percentage of specific lysis was calculated as the ratio of background deducted bioluminescence signals between cocultured and total lysed samples.

### Assessment of T cell exhaustion

T cells were activated by anti-hCD3 and anti-hCD28 (BioLegend, 317326, 302933) for two days, transduced with CAR retroviral plasmids, and cultured in the presence of 100 U/ml IL-2. On day 10 or 20, cells were stained with anti-PD1-FITC, anti-LAG3-APC, and anti-TIM3-BV510 (BioLegend, 329903, 369211, 345029), respectively, at 4 °C in dark for 30 min, washed, and analyzed by flow cytometry. Non-transduced cells were used as control.

## AUTHOR CONTRIBUTIONS

Z.W., A.S. and X.G. designed the research; Z.W. and A.S. performed experiments; Z.W., A.S., and X.G. analyzed and interpreted the data; Z.W., A.S. and X.G. wrote the manuscript; and X.G. supervised the study.

## ACKNOWLEDGEMENTS

This work was supported by R35GM141089. We thank NIH Tetramer Core Facility (contract number 75N93020D00005) for providing NY-ESO-1_157-165_/A2 pMHC-AF647 and MART1(A2L)_26-35_/A2 pMHC-AF647 tetramers.

## REFERENCES

Akatsuka Y (2020). TCR-Like CAR-T Cells Targeting MHC-Bound Minor Histocompatibility Antigens. Front Immunol. 11:257.

Allen F, Crepaldi L, Alsinet C, Strong AJ, Kleshchevnikov V, De Angeli P, Páleníková P, Khodak A, Kiselev V, Kosicki M, Bassett AR, Harding H, Galanty Y, Muñoz-Martínez F, Metzakopian E, Jackson SP, Parts L (2018). Predicting the mutations generated by repair of Cas9-induced double-strand breaks. Nat Biotechnol. 37: 64–72.

Andreatta M, Nielsen M (2016). Gapped sequence alignment using artificial neural networks: application to the MHC class I system. Bioinformatics. 32:511–517.

Banik D, Hamidinia M, Brzostek J, Wu L, Stephens HM, MacAry PA, Reinherz EL, Gascoigne NRJ, Lang MJ (2021). Single Molecule Force Spectroscopy Reveals Distinctions in Key Biophysical Parameters of αβ T-Cell Receptors Compared with Chimeric Antigen Receptors Directed at the Same Ligand. J Phys Chem Lett. 12:7566–7573.

Baulu E, Gardet C, Chuvin N, Depil S (2023). TCR-engineered T cell therapy in solid tumors: State of the art and perspectives. Sci Adv. 9:eadf3700.

Bendle GM, Linnemann C, Hooijkaas AI, Bies L, de Witte MA, Jorritsma A, Kaiser AD, Pouw N, Debets R, Kieback E, Uckert W, Song JY, Haanen JB, Schumacher TN (2010). Lethal graft-versus-host disease in mouse models of T cell receptor gene therapy. Nat Med. 16:565–570.

Bethune MT, Gee MH, Bunse M, Lee MS, Gschweng EH, Pagadala MS, Zhou J, Cheng D, Heath JR, Kohn DB, Kuhns MS, Uckert W, Baltimore D (2016). Domain-swapped T cell receptors improve the safety of TCR gene therapy. eLife. 5:e19095.

Boel A, De Saffel H, Steyaert W, Callewaert B, De Paepe A, Coucke PJ, Willaert A (2018). CRISPR/Cas9-mediated homology-directed repair by ssODNs in zebrafish induces complex mutational patterns resulting from genomic integration of repair-template fragments. Dis Model Mech. 11:dmm035352.

Bunse M, Bendle GM, Linnemann C, Bies L, Schulz S, Schumacher TN, Uckert W (2014). RNAi-mediated TCR knockdown prevents autoimmunity in mice caused by mixed TCR dimers following TCR gene transfer. Mol Ther. 22:1983–1991.

Cao J, Novoa EM, Zhang Z, Chen WCW, Liu D, Choi GCG, Wong ASL, Wehrspaun C, Kellis M, Lu TK (2021). High-throughput 5’ UTR engineering for enhanced protein production in non-viral gene therapies. Nat Commun. 12:4138.

Ceja MA, Khericha M, Harris CM, Puig-Saus C, Chen YY (2024). CAR-T cell manufacturing: Major process parameters and next-generation strategies. J Exp Med. 221:e20230903.

Chen C, Wang Z, Kang M, Lee KB, Ge X (2023). High-fidelity large-diversity monoclonal mammalian cell libraries by cell cycle arrested recombinase-mediated cassette exchange. Nucleic Acids Res. 51:e113.

Chen J, Qiu S, Li W, Wang K, Zhang Y, Yang H, Liu B, Li G, Li L, Chen M, Lan J, Niu J, He P, Cheng L, Fan G, Liu X, Song X, Xu C, Wu H, Wang H (2023). Tuning charge density of chimeric antigen receptor optimizes tonic signaling and CAR-T cell fitness. Cell Res. 33:341–354.

Choi HK, Cong P, Ge C, Natarajan A, Liu B, Zhang Y, Li K, Rushdi MN, Chen W, Lou J, Krogsgaard M, Zhu C (2023). Catch bond models may explain how force amplifies TCR signaling and antigen discrimination. Nat Commun. 14:2616.

Clement M, Knezevic L, Dockree T, McLaren JE, Ladell K, Miners KL, Llewellyn-Lacey S, Rubina A, Francis O, Cole DK, Sewell AK, Bridgeman JS, Price DA, van den Berg HA, Wooldridge L (2021). CD8 coreceptor-mediated focusing can reorder the agonist hierarchy of peptide ligands recognized via the T cell receptor. Proc Natl Acad Sci U S A. 118:e2019639118.

Dao T, Mun SS, Molvi Z, Korontsvit T, Klatt MG, Khan AG, Nyakatura EK, Pohl MA, White TE, Balderes PJ, Lorenz IC, O’Reilly RJ, Scheinberg DA (2022). A TCR mimic monoclonal antibody reactive with the "public" phospho-neoantigen pIRS2/HLA-A*02:01 complex. JCI Insight. 7:e151624.

Das DK, Feng Y, Mallis RJ, Li X, Keskin DB, Hussey RE, Brady SK, Wang JH, Wagner G, Reinherz EL, Lang MJ (2015). Force-dependent transition in the T-cell receptor β-subunit allosterically regulates peptide discrimination and pMHC bond lifetime. Proc Natl Acad Sci U S A. 112:1517–1522.

Dash P, Fiore-Gartland AJ, Hertz T, Wang GC, Sharma S, Souquette A, Crawford JC, Clemens EB, Nguyen THO, Kedzierska K, La Gruta NL, Bradley P, Thomas PG (2017). Quantifiable predictive features define epitope-specific T cell receptor repertoires. Nature. 547:89–93.

Di Roberto RB, Castellanos-Rueda R, Frey S, Egli D, Vazquez-Lombardi R, Kapetanovic E, Kucharczyk J, Reddy ST (2020). A Functional Screening Strategy for Engineering Chimeric Antigen Receptors with Reduced On-Target, Off-Tumor Activation. Mol Ther. 28:2564–2576.

Douglass J, Hsiue EH, Mog BJ, Hwang MS, DiNapoli SR, Pearlman AH, Miller MS, Wright KM, Azurmendi PA, Wang Q, Paul S, Schaefer A, Skora AD, Molin MD, Konig MF, Liu Q, Watson E, Li Y, Murphy MB, Pardoll DM, Bettegowda C, Papadopoulos N, Gabelli SB, Kinzler KW, Vogelstein B, Zhou S (2021). Bispecific antibodies targeting mutant RAS neoantigens. Sci Immunol. 6:eabd5515.

Duan Z, Li D, Li N, Lin S, Ren H, Hong J, Hinrichs CS, Ho M (2024). CAR-T cells based on a TCR mimic nanobody targeting HPV16 E6 exhibit antitumor activity against cervical cancer. Mol Ther Oncol. 32:200892.

Durrant MG, Fanton A, Tycko J, Hinks M, Chandrasekaran SS, Perry NT, Schaepe J, Du PP, Lotfy P, Bassik MC, Bintu L, Bhatt AS, Hsu PD (2023). Systematic discovery of recombinases for efficient integration of large DNA sequences into the human genome. Nat Biotechnol. 41:488–499.

Fahad AS, Chung CY, Lopez Acevedo SN, Boyle N, Madan B, Gutiérrez-González MF, Matus-Nicodemos R, Laflin AD, Ladi RR, Zhou J, Wolfe J, Llewellyn-Lacey S, Koup RA, Douek DC, Balfour HH Jr, Price DA, DeKosky BJ (2022). Immortalization and functional screening of natively paired human T cell receptor repertoires. Protein Eng Des Sel. 35:gzab034.

Fenis A, Demaria O, Gauthier L, Vivier E, Narni-Mancinelli E (2024). New immune cell engagers for cancer immunotherapy. Nat Rev Immunol. 24:471–486.

Govers C, Sebestyén Z, Coccoris M, Willemsen RA, Debets R (2010). T cell receptor gene therapy: strategies for optimizing transgenic TCR pairing. Trends Mol Med. 16:77–87.

Hebeisen M, Schmidt J, Guillaume P, Baumgaertner P, Speiser DE, Luescher I, Rufer N (2015). Identification of Rare High-Avidity, Tumor-Reactive CD8+ T Cells by Monomeric TCR-Ligand Off-Rates Measurements on Living Cells. Cancer Res. 75:1983–1991.

Holland CJ, Crean RM, Pentier JM, de Wet B, Lloyd A, Srikannathasan V, Lissin N, Lloyd KA, Blicher TH, Conroy PJ, Hock M, Pengelly RJ, Spinner TE, Cameron B, Potter EA, Jeyanthan A, Molloy PE, Sami M, Aleksic M, Liddy N, Robinson RA, Harper S, Lepore M, Pudney CR, van der Kamp MW, Rizkallah PJ, Jakobsen BK, Vuidepot A, Cole DK (2020). Specificity of bispecific T cell receptors and antibodies targeting peptide-HLA. J Clin Invest. 130:2673–2688.

Hsiue EH, Wright KM, Douglass J, Hwang MS, Mog BJ, Pearlman AH, Paul S, DiNapoli SR, Konig MF, Wang Q, Schaefer A, Miller MS, Skora AD, Azurmendi PA, Murphy MB, Liu Q, Watson E, Li Y, Pardoll DM, Bettegowda C, Papadopoulos N, Kinzler KW, Vogelstein B, Gabelli SB, Zhou S (2021). Targeting a neoantigen derived from a common TP53 mutation. Science. 371:eabc8697.

Joglekar AV, Leonard MT, Jeppson JD, Swift M, Li G, Wong S, Peng S, Zaretsky JM, Heath JR, Ribas A, Bethune MT, Baltimore D (2019). T cell antigen discovery via signaling and antigen-presenting bifunctional receptors. Nat Methods. 16:191–198.

Klatt MG, Dao T, Yang Z, Liu J, Mun SS, Dacek MM, Luo H, Gardner TJ, Bourne C, Peraro L, Aretz ZEH, Korontsvit T, Lau M, Kharas MG, Liu C, Scheinberg DA (2022). A TCR mimic CAR T cell specific for NDC80 is broadly reactive with solid tumors and hematologic malignancies. Blood. 140:861–874.

Klebanoff CA, Chandran SS, Baker BM, Quezada SA, Ribas A (2023). T cell receptor therapeutics: immunological targeting of the intracellular cancer proteome. Nat Rev Drug Discov. 22:996–1017.

Ku Z, Xie X, Davidson E, Ye X, Su H, Menachery VD, Li Y, Yuan Z, Zhang X, Muruato AE, I Escuer AG, Tyrell B, Doolan K, Doranz BJ, Wrapp D, Bates PF, McLellan JS, Weiss SR, Zhang N, Shi PY, An Z (2021). Molecular determinants and mechanism for antibody cocktail preventing SARS-CoV-2 escape. Nat Commun. 12:469.

Li AW, Lim WA (2020). Engineering cytokines and cytokine circuits. Science. 370:1034–1035.

Li Y, Jiang W, Mellins ED (2022). TCR-like antibodies targeting autoantigen-mhc complexes: a mini-review. Front Immunol. 13:968432.

Linette GP, Stadtmauer EA, Maus MV, Rapoport AP, Levine BL, Emery L, Litzky L, Bagg A, Carreno BM, Cimino PJ, Binder-Scholl GK, Smethurst DP, Gerry AB, Pumphrey NJ, Bennett AD, Brewer JE, Dukes J, Harper J, Tayton-Martin HK, Jakobsen BK, Hassan NJ, Kalos M, June CH (2013). Cardiovascular toxicity and titin cross-reactivity of affinity-enhanced T cells in myeloma and melanoma. Blood. 122:863–871.

Liu X, Xu Y, Xiong W, Yin B, Huang Y, Chu J, Xing C, Qian C, Du Y, Duan T, Wang HY, Zhang N, Yu JS, An Z, Wang R (2022). Development of a TCR-like antibody and chimeric antigen receptor against NY-ESO-1/HLA-A2 for cancer immunotherapy. J Immunother Cancer. 10:e004035.

Long AH, Haso WM, Shern JF, Wanhainen KM, Murgai M, Ingaramo M, Smith JP, Walker AJ, Kohler ME, Venkateshwara VR, Kaplan RN, Patterson GH, Fry TJ, Orentas RJ, Mackall CL (2015). 4-1BB costimulation ameliorates T cell exhaustion induced by tonic signaling of chimeric antigen receptors. Nat Med. 21:581–590.

Maricque BB, Chaudhari HG, Cohen BA (2019). A massively parallel reporter assay dissects the influence of chromatin structure on cis-regulatory activity. Nat Biotechnol. 37:90–95.

Martin AD, Wang X, Sandberg ML, Negri KR, Wu ML, Toledo Warshaviak D, Gabrelow GB, McElvain ME, Lee B, Daris ME, Xu H, Kamb A (2021). Re-examination of MAGE-A3 as a T-cell Therapeutic Target. J Immunother. 44:95–105.

Maus MV, Plotkin J, Jakka G, Stewart-Jones G, Rivière I, Merghoub T, Wolchok J, Renner C, Sadelain M (2016). An MHC-restricted antibody-based chimeric antigen receptor requires TCR-like affinity to maintain antigen specificity. Mol Ther Oncolytics. 3:1–9.

Moravec Z, Zhao Y, Voogd R, Cook DR, Kinrot S, Capra B, Yang H, Raud B, Ou J, Xuan J, Wei T, Ren L, Hu D, Wang J, Haanen JBAG, Schumacher TN, Chen X, Porter E, Scheper W (2024). Discovery of tumor-reactive T cell receptors by massively parallel library synthesis and screening. Nat Biotechnol. doi: 10.1038/s41587-024-02210-6. Epub ahead of print.

Morgan RA, Chinnasamy N, Abate-Daga D, Gros A, Robbins PF, Zheng Z, Dudley ME, Feldman SA, Yang JC, Sherry RM, Phan GQ, Hughes MS, Kammula US, Miller AD, Hessman CJ, Stewart AA, Restifo NP, Quezado MM, Alimchandani M, Rosenberg AZ, Nath A, Wang T, Bielekova B, Wuest SC, Akula N, McMahon FJ, Wilde S, Mosetter B, Schendel DJ, Laurencot CM, Rosenberg SA (2013). Cancer regression and neurological toxicity following anti-MAGE-A3 TCR gene therapy. J Immunother. 36:133–151.

Parthiban K, Perera RL, Sattar M, Huang Y, Mayle S, Masters E, Griffiths D, Surade S, Leah R, Dyson MR, McCafferty J (2019). A comprehensive search of functional sequence space using large mammalian display libraries created by gene editing. mAbs. 11:884–898.

Peri A, Salomon N, Wolf Y, Kreiter S, Diken M, Samuels Y (2023). The landscape of T cell antigens for cancer immunotherapy. Nat Cancer. 4:937–954.

Poorebrahim M, Mohammadkhani N, Mahmoudi R, Gholizadeh M, Fakhr E, Cid-Arregui A (2021). TCR-like CARs and TCR-CARs targeting neoepitopes: an emerging potential. Cancer Gene Ther. 28:581–589.

Raskin S, Van Pelt S, Toner K, Balakrishnan PB, Dave H, Bollard CM, Yvon E (2021). Novel TCR-like CAR-T cells targeting an HLA∗0201-restricted SSX2 epitope display strong activity against acute myeloid leukemia. Mol Ther Methods Clin Dev. 23:296–306.

Rydzek J, Nerreter T, Peng H, Jutz S, Leitner J, Steinberger P, Einsele H, Rader C, Hudecek M (2019). Chimeric Antigen Receptor Library Screening Using a Novel NF-κB/NFAT Reporter Cell Platform. Mol Ther. 27:287–299.

Salter AI, Rajan A, Kennedy JJ, Ivey RG, Shelby SA, Leung I, Templeton ML, Muhunthan V, Voillet V, Sommermeyer D, Whiteaker JR, Gottardo R, Veatch SL, Paulovich AG, Riddell SR (2021). Comparative analysis of TCR and CAR signaling informs CAR designs with superior antigen sensitivity and in vivo function. Sci Signal. 14:eabe2606.

Segaliny AI, Jayaraman J, Chen X, Chong J, Luxon R, Fung A, Fu Q, Jiang X, Rivera R, Ma X, Ren C, Zimak J, Hedde PN, Shang Y, Wu G, Zhao W. A high throughput bispecific antibody discovery pipeline. Commun Biol. (2023) 6:380.

Sengupta S, Board NL, Wu F, Moskovljevic M, Douglass J, Zhang J, Reinhold BR, Duke-Cohan J, Yu J, Reed MC, Tabdili Y, Azurmendi A, Fray EJ, Zhang H, Hsiue EH, Jenike K, Ho YC, Gabelli SB, Kinzler KW, Vogelstein B, Zhou S, Siliciano JD, Sadegh-Nasseri S, Reinherz EL, Siliciano RF (2022). TCR-mimic bispecific antibodies to target the HIV-1 reservoir. Proc Natl Acad Sci U S A. 119:e2123406119.

Sibener LV, Fernandes RA, Kolawole EM, Carbone CB, Liu F, McAffee D, Birnbaum ME, Yang X, Su LF, Yu W, Dong S, Gee MH, Jude KM, Davis MM, Groves JT, Goddard WA III, Heath JR, Evavold BD, Vale RD, Garcia KC (2018). Isolation of a Structural Mechanism for Uncoupling T Cell Receptor Signaling from Peptide-MHC Binding. Cell. 174:672–687.

Singh N, Frey NV, Engels B, Barrett DM, Shestova O, Ravikumar P, Cummins KD, Lee YG, Pajarillo R, Chun I, Shyu A, Highfill SL, Price A, Zhao L, Peng L, Granda B, Ramones M, Lu XM, Christian DA, Perazzelli J, Lacey SF, Roy NH, Burkhardt JK, Colomb F, Damra M, Abdel-Mohsen M, Liu T, Liu D, Standley DM, Young RM, Brogdon JL, Grupp SA, June CH, Maude SL, Gill S, Ruella M (2021). Antigen-independent activation enhances the efficacy of 4-1BB-costimulated CD22 CAR T cells. Nat Med. 27:842–850.

Spindler MJ, Nelson AL, Wagner EK, Oppermans N, Bridgeman JS, Heather JM, Adler AS, Asensio MA, Edgar RC, Lim YW, Meyer EH, Hawkins RE, Cobbold M, Johnson DS. (2020). Massively parallel interrogation and mining of natively paired human TCRαβ repertoires. Nat Biotechnol. 38:609–619.

Stewart-Jones G, Wadle A, Hombach A, Shenderov E, Held G, Fischer E, Kleber S, Nuber N, Stenner-Liewen F, Bauer S, McMichael A, Knuth A, Abken H, Hombach AA, Cerundolo V, Jones EY, Renner C (2009). Rational development of high-affinity T-cell receptor-like antibodies. Proc Natl Acad Sci U S A. 106:5784–5788.

Sockolosky JT, Trotta E, Parisi G, Picton L, Su LL, Le AC, Chhabra A, Silveria SL, George BM, King IC, Tiffany MR, Jude K, Sibener LV, Baker D, Shizuru JA, Ribas A, Bluestone JA, Garcia KC (2018). Selective targeting of engineered T cells using orthogonal IL-2 cytokine-receptor complexes. Science. 359:1037–1042.

Tousley AM, Rotiroti MC, Labanieh L, Rysavy LW, Kim WJ, Lareau C, Sotillo E, Weber EW, Rietberg SP, Dalton GN, Yin Y, Klysz D, Xu P, de la Serna EL, Dunn AR, Satpathy AT, Mackall CL, Majzner RG (2023). Co-opting signalling molecules enables logic-gated control of CAR T cells. Nature. 615:507–516.

van Loenen MM, de Boer R, Amir AL, Hagedoorn RS, Volbeda GL, Willemze R, van Rood JJ, Falkenburg JH, Heemskerk MH (2010). Mixed T cell receptor dimers harbor potentially harmful neoreactivity. Proc Natl Acad Sci. 107:10972–109777.

Uenishi GI, Repic M, Yam JY, Landuyt A, Saikumar-Lakshmi P, Guo T, Zarin P, Sassone-Corsi M, Chicoine A, Kellogg H, Hunt M, Drow T, Tewari R, Cook PJ, Yang SJ, Cerosaletti K, Schweinoch D, Guiastrennec B, James E, Patel C, Chen TF, Buckner JH, Rawlings DJ, Wickham TJ, Mueller KT (2024). GNTI-122: an autologous antigen-specific engineered Treg cell therapy for type 1 diabetes. JCI Insight. 9:e171844.

Varela-Rohena A, Molloy PE, Dunn SM, Li Y, Suhoski MM, Carroll RG, Milicic A, Mahon T, Sutton DH, Laugel B, Moysey R, Cameron BJ, Vuidepot A, Purbhoo MA, Cole DK, Phillips RE, June CH, Jakobsen BK, Sewell AK, Riley JL (2008). Control of HIV-1 immune escape by CD8 T cells expressing enhanced T-cell receptor. Nat Med. 14:1390–1395.

Vazquez-Lombardi R, Jung JS, Schlatter FS, Mei A, Mantuano NR, Bieberich F, Hong KL, Kucharczyk J, Kapetanovic E, Aznauryan E, Weber CR, Zippelius A, Läubli H, Reddy ST (2022). High-throughput T cell receptor engineering by functional screening identifies candidates with enhanced potency and specificity. Immunity. 55:1953–1966.

Wang JH (2020). T cell receptors, mechanosensors, catch bonds and immunotherapy. Prog Biophys Mol Biol. 153:23–27.

Wang L, Matsumoto M, Akahori Y, Seo N, Shirakura K, Kato T, Katsumoto Y, Miyahara Y, Shiku H (2024). Preclinical evaluation of a novel CAR-T therapy utilizing a scFv antibody highly specific to MAGE-A4_p230-239_/HLA-A∗02:01 complex. Mol Ther. 32:734–748.

Wang X, Martin AD, Negri KR, McElvain ME, Oh J, Wu ML, Lee WH, Ando Y, Gabrelow GB, Toledo Warshaviak D, Sandberg ML, Xu H, Kamb A (2021). Extensive functional comparisons between chimeric antigen receptors and T cell receptors highlight fundamental similarities. Mol Immunol. 138:137–149.

Wang Z, Chen C, Ge X (2024a). Large T antigen mediated target gene replication improves site-specific recombination efficiency. Front Bioeng Biotechnol. 12:1377167.

Wang Z, Kang M, Ebrahimpour A, Chen C, Ge X (2024b). Fc engineering by monoclonal mammalian cell display for improved affinity and selectivity towards FcγRs. Antib Ther. 7:209–220.

Webb AI, Dunstone MA, Chen W, Aguilar MI, Chen Q, Jackson H, Chang L, Kjer-Nielsen L, Beddoe T, McCluskey J, Rossjohn J, Purcell AW (2004). Functional and structural characteristics of NY-ESO-1-related HLA A2-restricted epitopes and the design of a novel immunogenic analogue. J Biol Chem. 279:23438–23446.

Yang X, Nishimiya D, Löchte S, Jude KM, Borowska M, Savvides CS, Dougan M, Su L, Zhao X, Piehler J, Garcia KC (2023). Facile repurposing of peptide-MHC-restricted antibodies for cancer immunotherapy. Nat Biotechnol. 41:932–943.

Yarmarkovich M, Marshall QF, Warrington JM, Premaratne R, Farrel A, Groff D, Li W, di Marco M, Runbeck E, Truong H, Toor JS, Tripathi S, Nguyen S, Shen H, Noel T, Church NL, Weiner A, Kendsersky N, Martinez D, Weisberg R, Christie M, Eisenlohr L, Bosse KR, Dimitrov DS, Stevanovic S, Sgourakis NG, Kiefel BR, Maris JM (2023). Targeting of intracellular oncoproteins with peptide-centric CARs. Nature. 623:820–827.

Zhang L, Sosinowski T, Cox AR, Cepeda JR, Sekhar NS, Hartig SM, Miao D, Yu L, Pietropaolo M, Davidson HW (2019). Chimeric antigen receptor (CAR) T cells targeting a pathogenic MHC class II:peptide complex modulate the progression of autoimmune diabetes. J Autoimmun. 96:50–58.

Zhao X, Kolawole EM, Chan W, Feng Y, Yang X, Gee MH, Jude KM, Sibener LV, Fordyce PM, Germain RN, Evavold BD, Garcia KC (2022). Tuning T cell receptor sensitivity through catch bond engineering. Science. 376:eabl5282.

Zhu C, Chen W, Lou J, Rittase W, Li K (2019). Mechanosensing through immunoreceptors. Nat Immunol. 20:1269–1278.

Zdinak PM, Trivedi N, Grebinoski S, Torrey J, Martinez EZ, Martinez S, Hicks L, Ranjan R, Makani VKK, Roland MM, Kublo L, Arshad S, Anderson MS, Vignali DAA, Joglekar AV (2024). De novo identification of CD4+ T cell epitopes. Nat Methods. 21:846–856.

