## Supplementary material for "De novo functional discovery of peptide-MHC restricted CARs from recombinase-constructed large-diversity monoclonal T cell libraries": Fig. S

**Contents:**

**Figure S1.** Recombinase-based transgene integration to polyclonal Jurkat cells.

**Figure S2.** Flow cytometry plots of selected monoclonal Jurkat-Pa01 cells.

**Figure S3.** Genotyping of Jurkat-Pa01 E11 and B4 clones.

**Figure S4.** Integration of transgene iRFP to Jurkat-Pa01 cell lines.

**Figure S5.** Integration of transgenes MA2 CAR to Jurkat-Pa01 B4 cells.

**Figure S6.** Germline usage of identified unique TCRm CARs targeting different antigens.

**Figure S7.** Representative NFAT-EGFP screening data.

**Figure S8.** Peptide sequences and affinities.

**Figure S9.** In vitro killing shown by fluorescence microscope imaging.

**Figure S10.** Exhaustion markers on day 20 post transduction.

**Figure S1:**

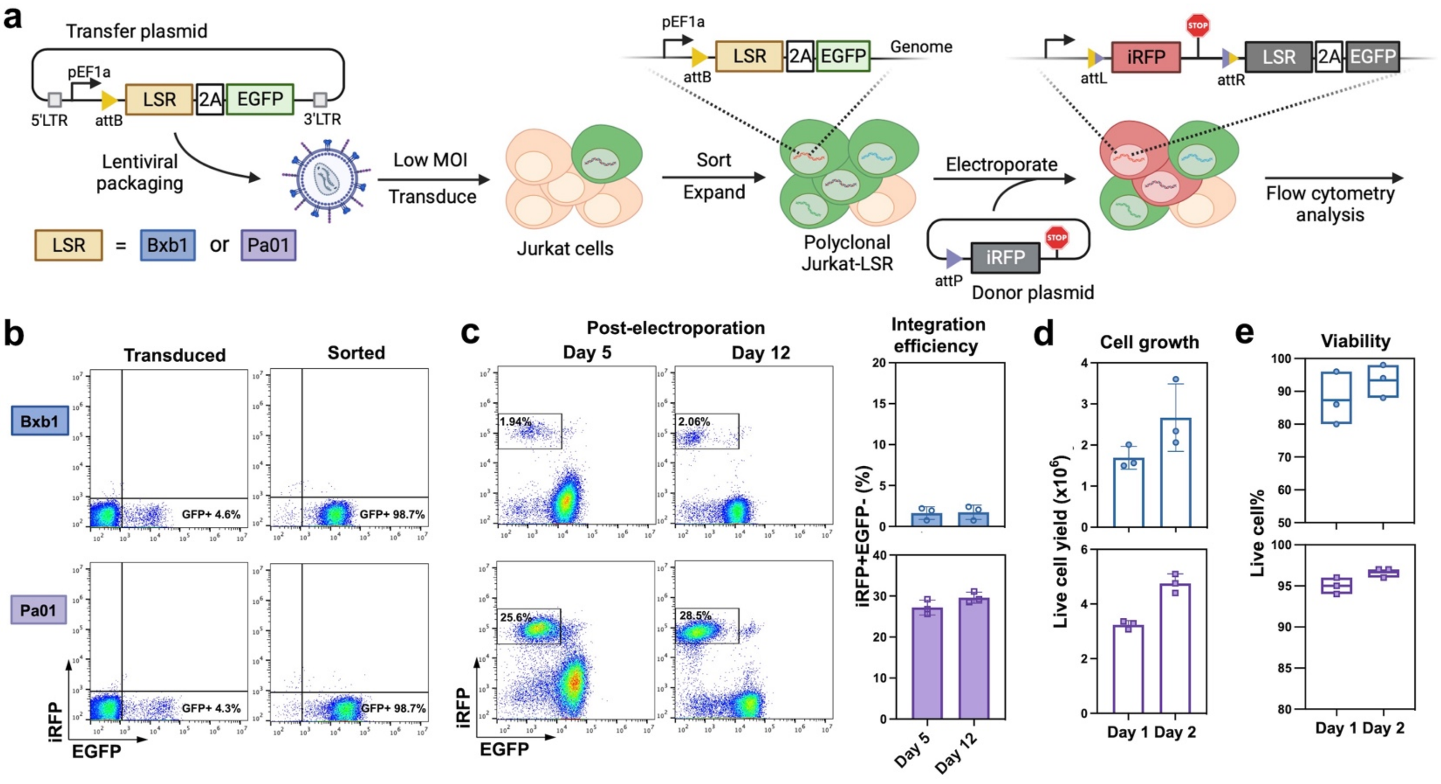


**Figure S1. Recombinase-based transgene integration to polyclonal Jurkat cells**. **a,** Schematic of generation of Jurkat cells carrying a landing pad on the genome and recombinase-based transgene integration. **b,** Flow cytometry plots showing Jurkat cells transduced with pEF1a-attB-LSR-T2A-EGFP landing pad cassette before sorting and after sorting. **c,** Comparison of BxB1 and Pa01 integration efficiency. Two million of polyclonal Jurkat-LSR cells were electroporated with 4 µg of corresponding donor plasmid attP-iRFP and analyzed by flow cytometry on day 5 and day 12 post electroporation. **d,** Cell growth of polyclonal Jurkat-LSR after electroporation with donor plasmid attP-iRFP. **e,** Cell viability of polyclonal Jurkat-LSR after electroporation with donor plasmid attP-iRFP.

**Figure S2:**

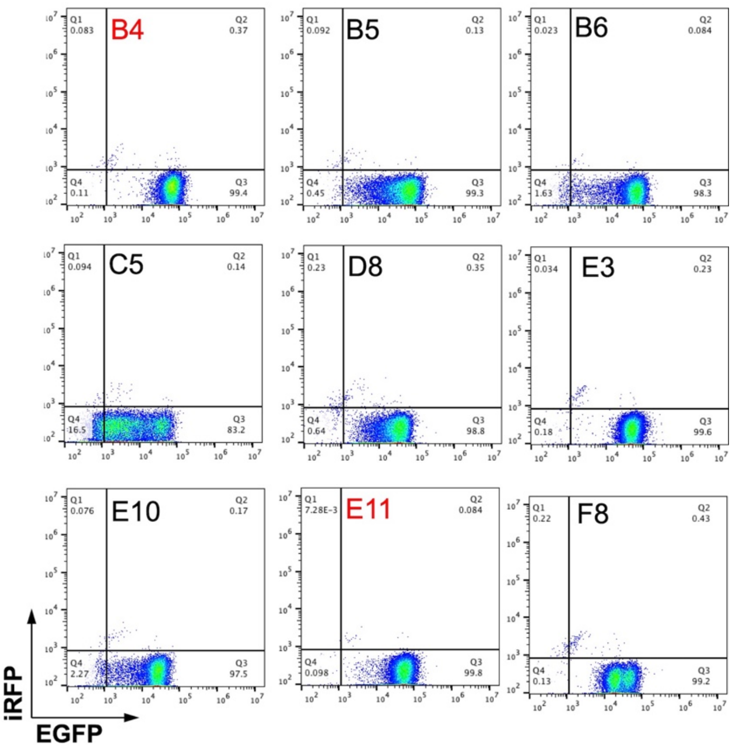


**Figure S2. Flow cytometry plots of selected monoclonal Jurkat-Pa01 cells.**

**Figure S3:**

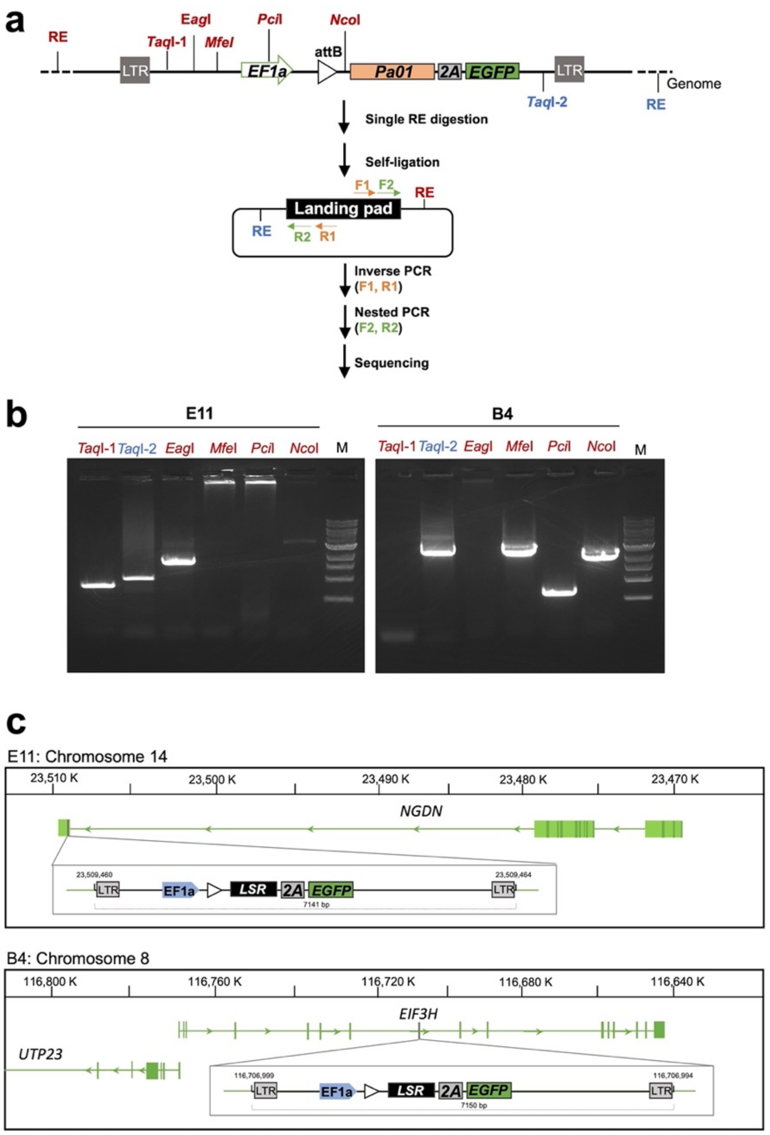


**Figure S3. Genotyping of Jurkat-Pa01 E11 and B4 clones. a,** Schematic of inverse PCR (iPCR) for landing pad genotyping. Genomic DNA was extracted and digested with one of the indicated restriction enzymes, self-ligated, and subjected to iPCRs in which primers were designed to outwardly anneal at the integrated landing region. **b,** Electrophoresis analysis of iPCR products. **c,** Genomic context of the landing pads of Jurkat-Pa01 E11 and B4 revealed by Sanger sequencing. Human genome sequence (GRCh38.p14) was used as the reference.

**Figure S4:**

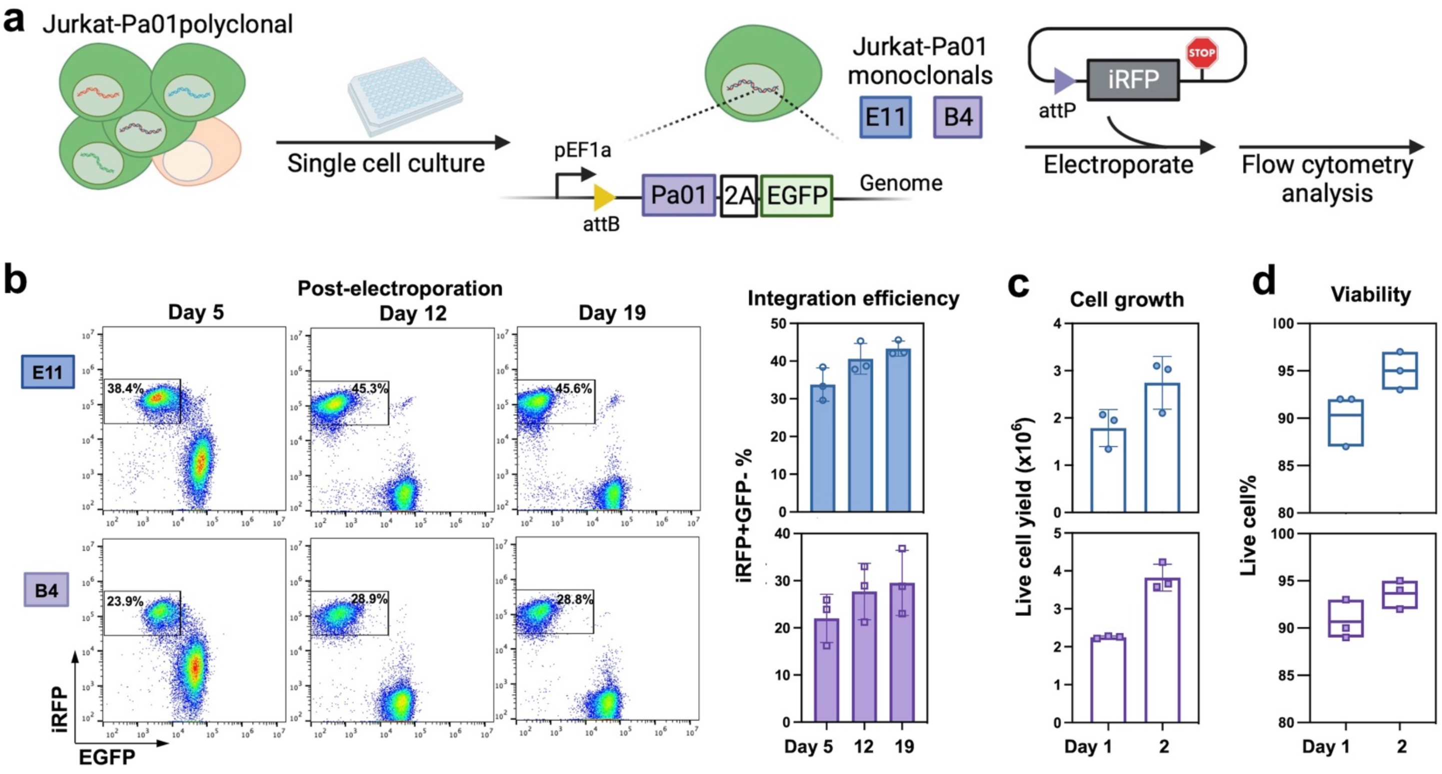


**Figure S4. Integration of transgene iRFP to Jurkat-Pa01 cell lines**. **a,** Schematic of monoclonal Jurkat-Pa01 cell selection and recombinase-based transgene integration. **b,** The integration efficiency in monoclonal Jurkat-Pa01 cell lines E11 and B4. Two million of monoclonal Jurkat-Pa01 cells were electroporated with 4 μg of donor plasmid attP-iRFP and analyzed by flow cytometry on day 5, 12, and 19 post electroporation. **c,** Cell growth of monoclonal Jurkat-Pa01 after electroporation with donor plasmid attP-iRFP. **d,** Cell viability of monoclonal Jurkat-Pa01 after electroporation with donor plasmid attP-iRFP.

**Figure S5:**

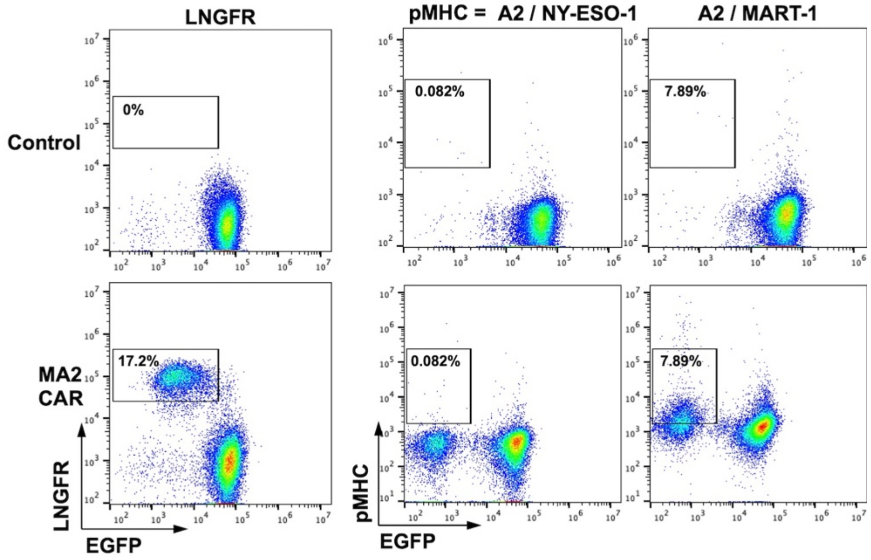


**Figure S5. Integration of transgenes MA2 CAR to Jurkat-Pa01 B4 cells.** Two million of monoclonal Jurkat-Pa01 B4 cells were electroporated with 4 μg of donor plasmid attP-MA2 and analyzed by flow cytometry on day 5. Cells without electroporation were used as control.

**Figure S6:**

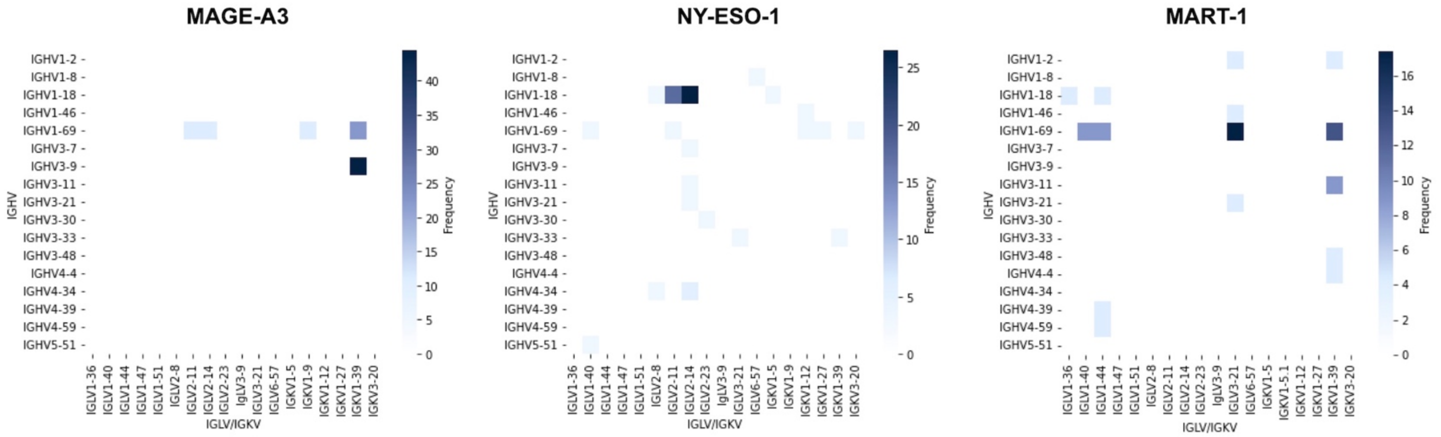


**Figure S6. Germline usage of identified unique TCRm CARs targeting different antigens.**

**Figure S7:**

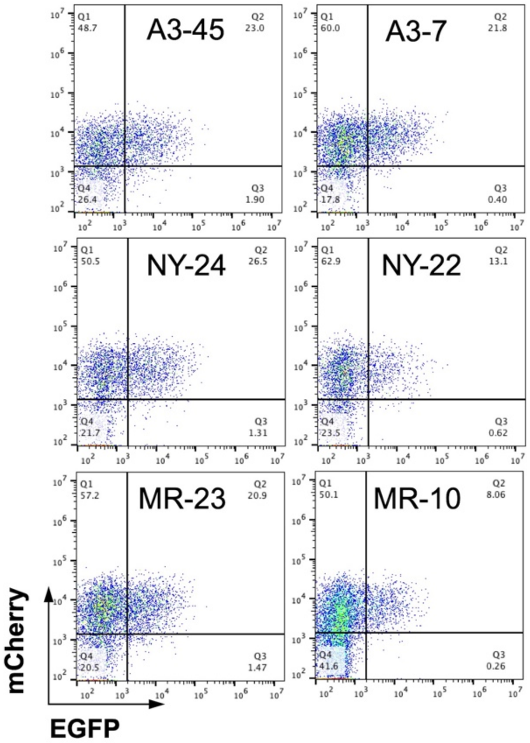


**Figure S7. Representative NFAT-EGFP screening data.** Jurkat NFAT-EGFP reporter cells were transduced with TCR-like CAR-2A-mCherry lentiviral plasmids, and overnight cocultured with cognate peptide pulsed 293F cells. Cell populations of CAR expression (mCherry$^+$, Q1+Q2) and activation (EGFP$^+$, Q2) were determined by flow cytometry. The relative percentages of EGFP$^+$ over mCherry$^+$ cells were calculated as the levels of T cell activation.

**Figure S8:**

| Gene | Peptide | Affinity to HLA-A*02:01 (nM) |
|---|---|---|
| **NY-ESO-1$_{157-164}$** | **SLLMWITQC** | **422.15** |
| CYBB$_{172-180}$ | TLLAGITGV | 5.49 |
| ABCC$_{35-43}$ | SLLAWVPCI | 5.85 |
| **MAGE-A3$_{112-120}$** | **KVAELVHFL** | **11.32** |
| EPS8L2$_{339-347}$ | SAAELVHFL | 267.44 |
| SLC40A1$_{501-509}$ | YLLDLLHFI | 1.84 |
| HK3$_{314-322}$ | YLGELVRLV | 7.41 |
| MAGE-A8$_{115-123}$ | KVAELVRFL | 195.91 |
| **MART-1$_{26-35(27L)}$** | **ELAGIGILTV** | **215.85** |
| LY6G6C$_{115-124}$ | SLAGLGLWLL | 13.89 |
| DNAJC4$_{164-173}$ | MLAGMGLHYI | 8.01 |

https://marisshiny.research.chop.edu/sCRAP
https://services.healthtech.dtu.dk/services/NetMHC-4.0

| Gene | Peptide | Affinity to HLA-A*02:01 (nM) |
|---|---|---|
| **NY-ESO-1$_{157-164}$** | **SLLMWITQC** | **422.15** |
| S1A | ALLMWITQC | 736.18 |
| L2A | SALMWITQC | 10449.59 |
| L3A | SLAMWITQC | 452.58 |
| M4A | SLLAWITQC | 102.1 |
| W5A | SLLMAITQC | 1059.93 |
| I6A | SLLMWATQC | 1361.57 |
| T7A | SLLMWIAQC | 390.92 |
| Q8A | SLLMWITAC | 483.55 |
| C9A | SLLMWITQA | 17.12 |
| **MAGE-A3$_{112-120}$** | **KVAELVHFL** | **11.32** |
| K1A | AVAELVHFL | 19.18 |
| V2A | KAAELVHFL | 154.98 |
| E4A | KVAALVHFL | 31.63 |
| L5A | KVAEAVHFL | 18.47 |
| V6A | KVAELAHFL | 29.43 |
| H7A | KVAELVAFL | 13.36 |
| F8A | KVAELVHAL | 27.5 |
| L9A | KVAELVHFA | 57.8 |

**Figure S8. Peptide sequences and affinities.** The cognate peptides are shown in bold, and top hits of potential cross-reactive peptides were predicated by sCRAP [Yarmarkovich *et al.*, 2023]. Affinities to HLA-A*02:01 were predicted by NetMHCpan 4.0 [Andreatta & Nielsen, 2016].

**Figure S9:**

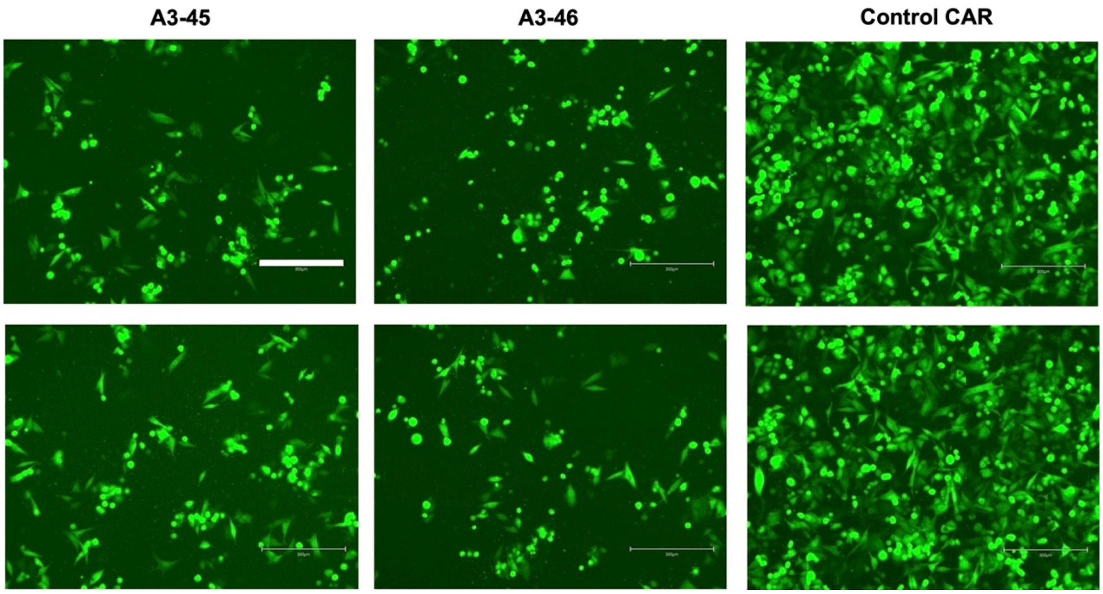


**Figure S9. In vitro killing shown by fluorescence microscope imaging.** MAGE-A3 CAR transduced primary T cells were cocultured with target A-375 cells overnight. Effector: Target = 5:1. Non-functional CAR A3-53 was used as control. Bar = 300 μm.

**Figure S10:**

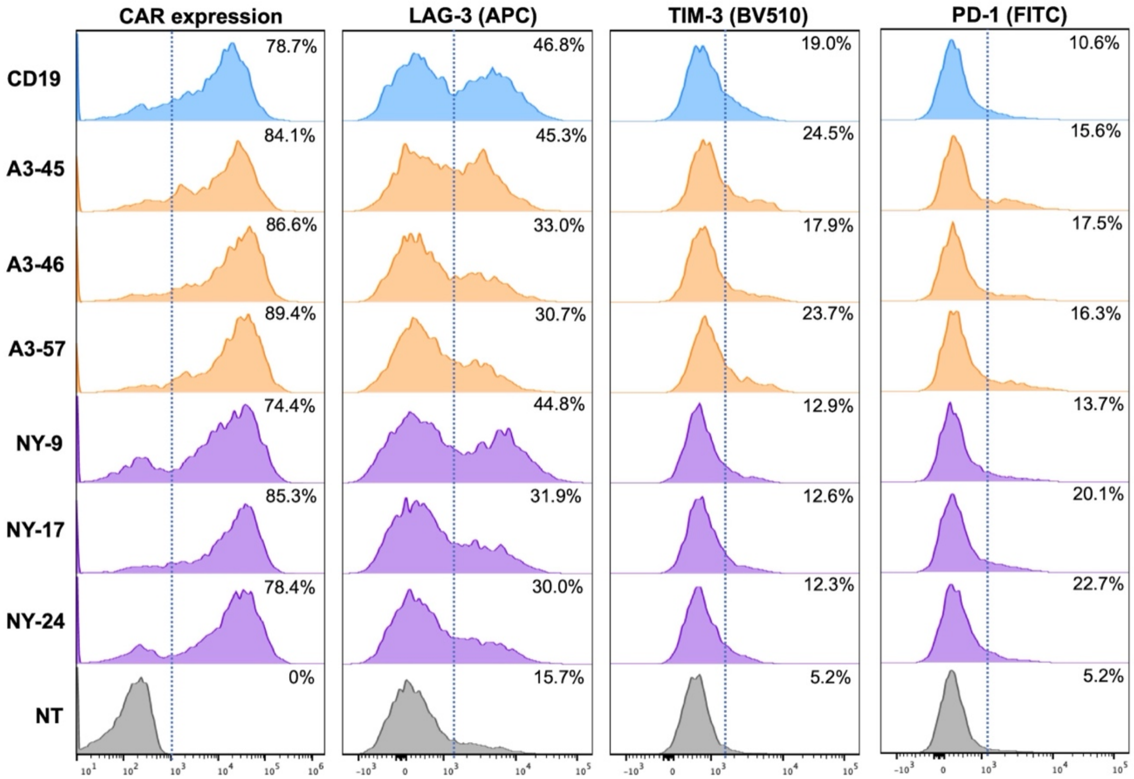


**Figure S10. Exhaustion markers on day 20 post transduction.** Flow cytometry histograms showing expression of CAR, LAG-3, TIM-3, and PD-1.
